# The many dimensions of combination therapy: How to combine antibiotics to limit resistance evolution

**DOI:** 10.1101/2023.11.29.569181

**Authors:** Christin Nyhoegen, Sebastian Bonhoeffer, Hildegard Uecker

## Abstract

In combination therapy, bacteria are challenged with two or more antibiotics simultaneously. Ideally, separate mutations are required to adapt to each of them, which is *a priori* expected to hinder the evolution of full resistance. Yet, the success of this strategy ultimately depends on how well the combination controls the growth of bacteria with and without resistance mutations. To design a combination treatment, we need to choose drugs and their doses and decide how many drugs get mixed. Which combinations are good? To answer this question, we set up a stochastic pharmacodynamic model and determine the probability to successfully eradicate a bacterial population. We consider bacteriostatic and two types of bactericidal drugs – those that kill independent of replication and those that kill during replication. To establish results for a null model, we consider non-interacting drugs and implement the two most common models for drug independence – Loewe additivity and Bliss independence. Our results show that combination therapy is almost always better in limiting the evolution of resistance than administering just one drug, even though we keep the total drug dose constant for a ‘fair’ comparison. Yet, exceptions exist for drugs with steep dose-response curves. Combining a bacteriostatic and a bactericidal drug which can kill non-replicating cells is particularly beneficial. Our results suggest that a 50:50 drug ratio – even if not always optimal – is usually a good and safe choice. Applying three or four drugs is beneficial for treatment of strains with large mutation rates but adding more drugs otherwise only provides a marginal benefit or even a disadvantage. By systematically addressing key elements of treatment design, our study provides a basis for future models which take further factors into account. It also highlights conceptual challenges with translating the traditional concepts of drug independence to the single-cell level.

## 1 Introduction

Antibiotic resistance poses a major challenge for patient treatment worldwide. The rapid evolution and spread of resistance results in a loss of treatment options, increasing the demand for new antibiotics. Designing treatment strategies that limit the evolution of resistance can potentially extend the lifetime of the antibiotics currently in use and therefore secure the availability of effective treatments for the future (Tyers & Wright, 2019). One possibility to reduce the risk of resistance evolution during treatment is to increase the genetic barrier to resistance, i.e. to increase the number of mutations (or resistance genes) the bacteria need to escape the treatment. This can especially be achieved by using more than one drug in combination. Ideally (i.e. in the absence of cross-resistance), bacteria then need to acquire multiple mutations – one for each drug – to become fully resistant as opposed to a single mutation, which would be sufficient for resistance to mono-therapy. The most well-known example of the efficiency of combination treatment of bacterial infections is the treatment of *Mycobacterium tuberculosis* infections, for which drug combinations have been the standard treatment for more than 70 years due to their ability to reduce resistance evolution (Medical Research Council, 1950; Fox et al., 1999). As drug resistance is becoming a problem for the treatment of more and more different bacterial infections, many researchers in evolutionary medicine highly recommend using combination therapy more widely (Bonhoeffer et al., 1997; Palmer & Kishony, 2013; Roemhild & Schulenburg, 2019; Andersson et al., 2020; Merker et al., 2020; Woods & Read, 2023).

In order to optimally design combination treatments, we need to gain a thorough understanding of how the choice of drugs and their precise administration affect the evolution of resistance. To prevent the accumulation of resistance mutations up to full resistance, a good combination treatment must limit the appearance of new mutations and the spread of partially resistant genotypes.

The efficiency of combinations in limiting resistance evolution depends on drug characteristics such as the bacteriostatic vs bactericidal activity (in the following referred to as modes of action), their dose-response curves, interactions between the drugs and collateral effects, and complex physiological responses such as hysteresis. Interestingly, the effects of drug-drug interactions and collateral effects – two rather complex factors – on resistance evolution are more frequently explored (Hegreness et al., 2008; Pena-Miller et al., 2013; Munck et al., 2014; Barbosa et al., 2018) than those of the modes of action and the shape of the dose-response curves, which seem more ‘fundamental’. For comprehensive reviews on the former, see, for example, Bollenbach (2015), Baym et al. (2016), and Roemhild et al. (2022). Several studies explored the effect of the drugs’ modes of action on the type of drug-drug interactions – synergism or antagonism – (Ocampo et al., 2014; Chandrasekaran et al., 2016; Lázár et al., 2022). Coates et al. (2018) further showed that the efficiency of a combination in clearing a susceptible population depends on the modes of action of the combined drugs. Yet, to our knowledge, a direct assessment of how the modes of action of drugs in combination influence resistance evolution is missing. Besides the mode of action, the shape of the dose-response curve is another factor to consider when combining drugs. A study by Russ and Kishony (2018) showed, for example, that drugs with steep dose-response curves need to be applied at a higher total drug concentration to inhibit the growth of a bacterial population to the same extent as mono-therapy.

For optimal combination therapy, we do not only need to know *which* drugs to apply but also *how*. With multiple drugs in combination, applying the different drugs, each at the same concentration as in mono-therapy, may not be necessary to clear the wild-type infection. In fact, lowering the doses might even be required to avoid toxicity. Increasing the number of drugs combined while keeping the total drug dose constant can reduce the drugs’ effect on the growth of the bacterial population (Russ & Kishony, 2018; Lázár et al., 2022). How such a reduction in individual doses would impair the efficiency of combination therapy in limiting resistance evolution is unclear, especially with a larger number of drugs. While the above studies split the total drug dose equally among the drugs, unequal drug ratios could be beneficial, especially when the drugs vary in their characteristics.

In this study, we set up a stochastic pharmacodynamic model to test the efficiencies of various combination treatments when resistance evolution relies on mutations in the chromosome. We compare treatments based on their probability of successfully eradicating a bacterial population, which we derive from branching process theory. Besides calculating the probability of treatment success, we also look at the two factors that determine the risk of treatment failure due to resistance evolution – the expected number of resistant mutants and their chance to escape stochastic loss while rare. This allows us to understand why one treatment works better than another. We focus on the effects of the modes of action, the dose-response relationships, the number of drugs and their doses and exclude drug-drug interactions and collateral effects. Calculating the combined effect of drugs in the absence of drug-drug interactions requires choosing a null model for drug additivity. In our analysis, we employ the two most common models, Bliss independence (Bliss, 1939) and Loewe additivity (Loewe & Muischnek, 1926). Bliss (1939) calls two drugs independent if the fraction of cells surviving combination treatment is the product of the single-drug survival probabilities. Loewe and Muischnek (1926) considers two drugs independent when a fraction of one drug can be exchanged by a fraction of the other without a change in drug effect. Our systematic analysis provides an overview over the influence of a range of treatment choices and allows us to formulate some rules of thumb for the design of promising treatments.

## 2 Methods

### 2.1 General model

We consider a bacterial population of initial size *N*_0_, undergoing treatment with *n* drugs. Bacteria can be fully susceptible wild-type cells (denoted by *W*) or resistant to any subset *I* of drugs (denoted by *M*_*I*_ with *I ⊆ {*1, 2, …, *n}*). A bacterium acquires a resistance mutation to drug *i* during replication with probability *u*_*i*_. We assume that each mutation confers resistance to exactly one drug, there is hence no cross resistance. Multiple mutations can appear simultaneously. The initial population is either fully susceptible, or resistant cells pre-exist at low frequency prior to treatment. Whenever we include standing genetic variation, we assume that the population is at mutation-selection balance (see appendix A). We ignore resource competition and other interactions between bacteria and assume that cells replicate and die independent of each other. The population dynamics are thus described by a multi-type branching process.

In the absence of treatment, wild-type bacteria replicate at an intrinsic rate 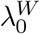 and die at an intrinsic rate *μ*_0_, resulting in a net *per capita* growth rate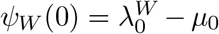. We assume that resistance to drug *i* entails a cost, reducing the replication rate by a factor (1*−γ*_*i*_), hence 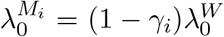 for a single mutant with resistance to drug *i*. The cost is multiplicative for types that are resistant to multiple drugs.

During treatment with drugs at concentration (*c*_1_, *c*_2_, *· · ·, c*_*n*_), the net *per capita* growth rate *ψ*_*X*_(*c*_1_, *c*_2_, *· · ·, c*_*n*_) of type *X* (with *X ∈ {W, M*_1_, *· · · }*) is reduced by a function *E*_*X*_(*c*_1_, *c*_2_, *· · ·, c*_*n*_), describing the effect of the antibiotic on the growth rate:

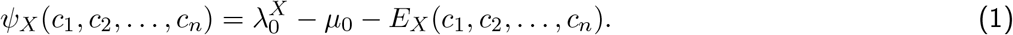

In the following sections, we provide derivations of the drug effect *E*_*X*_(*c*_1_, *c*_2_, *· · ·, c*_*n*_) and how it results from drug effects on replication and death rates for different modes of action.

### 2.2 Effect of mono-therapy for different modes of action

We describe the net *per-capita* growth rate in the presence of treatment with a single drug (*n* = 1) by a sigmoidal Hill function using the formulation by Regoes et al. (2004):

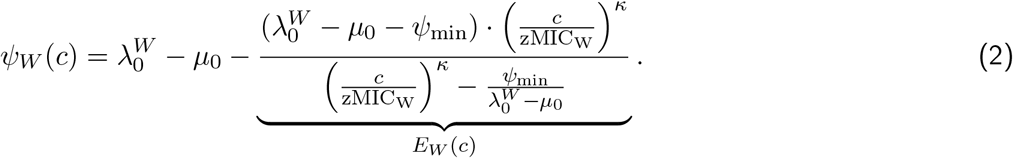

This dose-response relationship describes the pharmacodynamics of the drug. The susceptibility parameter zMIC_*W*_ is the concentration at which the net growth rate is zero. In the limit of very high concentrations, the net growth rate converges to *ψ*_min_ (lim_*c→∞*_ *ψ*(*c*) = *ψ*_min_ with *ψ*_min_ *<* 0*h*^*−*1^). The Hill coefficient *κ* describes the steepness of the curve.

We assume that the Hill coefficient *κ* and the parameter *ψ*_min_ are the same for all cell types, which can be observed in experimentally measured dose-response curves (Chevereau et al., 2015; Das et al., 2020), but might not to be universally true. The cost of resistance affects the intrinsic replication rate, as discussed earlier, and resistance provides a benefit of increasing the wild-type parameter zMIC_W_ by a factor *β*_res_. The net growth rate of a resistant mutant is thus calculated by

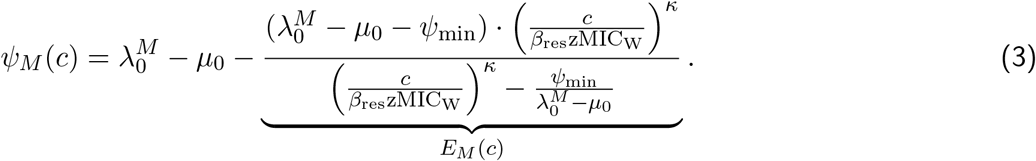

The drug effect on the net growth rate can be due to an increase in the death rate, a decrease in the replication rate or both. We refer to this as the ‘mode of action’ of a drug. Note that according to our definition, ‘mode of action’ thus does not relate to the specific mechanism of action or target site. For simplicity, we will drop the index indicating the cell type when describing the modes of action.

#### Bactericidal drug that increases the kill rate independent of replication (CK)

The first mode of action that we consider are bactericidal drugs (indicated by ‘CK’) that increase the kill rate independent of replication and have no effect on the replication rate. An example for this drug type are aminoglycoside antibiotics, which are bactericidal and able to kill non-replicating cells (Mutschler et al., 2005; McCall et al., 2019). We thus have *λ*^CK^(*c*) = *λ*_0_ and *μ*^CK^(*c*) = *μ*_0_ + *η*(*c*), where the function *η*(*c*) describes the drug-induced killing. Comparing the net growth rate

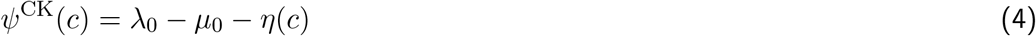

to equation (1), we see that *η*(*c*) = *E*(*c*) with *E*(*c*) given in equation (2) or (3). We thus have *μ*^CK^(*c*) = *μ*_0_ + *E*(*c*).

#### Bacteriostatic drug that reduces the replication rate (S)

At the other end of the spectrum, we consider bacteriostatic drugs, such as chloramphenicol, tetracycline or macrolide antbiotics (Mutschler et al., 2005), that only affect the replication but not the death rate (indicated by ‘S’). The replication rate is reduced by a factor (1 *− σ*(*c*)), hence *λ*^S^(*c*) = (1 *− σ*(*c*))*λ*_0_ (with *σ*(*c*) *∈* [0, 1] *∀c*). The net growth rate is given by

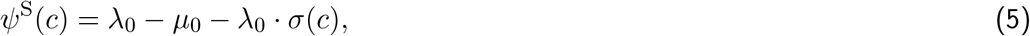

from which we can read off that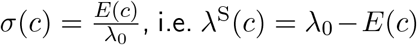, i.e. *λ*^S^(*c*) = *λ*_0_ *− E*(*c*). For this mode of action, the net growth rate at high concentrations, *ψ*_min_, can be at most *−μ*_0_, since the function *σ*(*c*) cannot be larger than one.

#### Bactericidal drug that acts on replicating cells (CR)

Many bactericidal drugs only act on metabolically active cells, such as *β*-lactam antibiotics, as well as drugs from other classes, for example ciprofloxacin or rifampicin (McCall et al., 2019). This is not captured by our CK drug, where killing is independent of the replication rate *λ*_0_ (which is correlated with the metabolic activity). In particular, some bactericidal drugs act during replication itself, and we include this as a third mode of action. We describe the fraction of cell replications which result in cell death by a function *π*(*c*) (with *π*(*c*) *∈* [0, 1] *∀c*). The replication rate is thus reduced to *λ*^CR^(*c*) = *λ*_0_ *·* (1 *− π*(*c*)) and the death rate increased to *μ*^CR^(*c*) = *μ*_0_ + *π*(*c*) *· λ*_0_, which gives the net growth rate

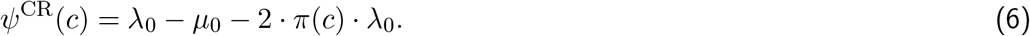

The function *π*(*c*) is thus given by 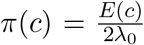. We therefore have 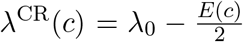 and 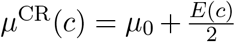. The lower limit of the parameter *ψ*_min_ is *−μ*_0_ *− λ*_0_, since *π*(*c*) *≤* 1.

Table 1 provides a summary of the net growth rates and replication and death rates for all three modes of action.

**Table 1:** Net growth rate, replication and death rate for the mono-therapy of drugs with different modes of action.

| Mode | $\psi(c)$ | $\lambda := \lambda(c)$ | $\mu := \mu(c)$ | Ref. eq. (2) |
| --- | --- | --- | --- | --- |
| CK | $\psi^{\text{CK}}(c) = \lambda_0 - \mu_0 - \eta(c)$ | $\lambda^{\text{CK}} = \lambda_0$ | $\mu^{\text{CK}} = \mu_0 + \eta(c)$ | $\eta(c) = E(c)$ |
| CR | $\psi^{\text{CR}}(c) = \lambda_0 - \mu_0 - 2\pi(c)\lambda_0$ | $\lambda^{\text{CR}} = \lambda_0 \cdot (1 - \pi(c))$ | $\mu^{\text{CR}} = \mu_0 + \lambda_0 \cdot \pi(c)$ | $\pi(c) = \frac{E(c)}{2\lambda_0}$ |
| S | $\psi^S(c) = \lambda_0 - \mu_0 - \sigma(c)\lambda_0$ | $\lambda^S = \lambda_0 \cdot (1 - \sigma(c))$ | $\mu^S = \mu_0$ | $\sigma(c) = \frac{E(c)}{\lambda_0}$ |

### 2.3 Modelling the combined effect of multiple antibiotics

Modelling the combined effect of multiple drugs requires choosing a null model for the effect of the drug combination in the absence of drug-drug interactions. The most common reference models are Bliss independence (Bliss, 1939) and Loewe additivity (Loewe & Muischnek, 1926; Loewe, 1928). Although the two approaches are often presented as ‘rival approaches’ (Greco et al., 1995), a glance at the original literature reveals that they are defined for different (complementary) situations – Bliss independence for drugs acting on different target sites and Loewe additivity for drugs with the same target. In the following, we will give a brief overview of the history of the two concepts, before we describe how we implement Bliss independence and Loewe additivity in our model.

#### 2.3.1 Background on the history of Bliss independence and Loewe additivity

Bliss independence and Loewe additivity were developed as reference models for ‘no interaction’ to determine whether a combination of drugs can be classified as synergistic or antago-nistic or, in other words, whether the combined effect of the drugs would be larger or smaller than expected under the assumption of zero interaction. However, answering how exactly the combined effect in the absence of interactions can be calculated is not trivial. Let us have a look at the original definitions of the additivity models.

Before discussing the seminal publications by Loewe and Bliss (Loewe & Muischnek, 1926; Loewe, 1928; Bliss, 1939), we want to highlight the work by Wilhelm Frei, published more than ten years before Loewe’s first paper on drug additivity (and, just as Loewe’s first paper, written in German). In 1913, Wilhelm Frei discusses, on a theoretical basis, how the joint effect of a drug combination could be predicted in order to compare the effect of combinations with mono-therapy for a fixed total dose, varying the ratios of the drugs in combination (Frei, 1913): Let us assume we use two drugs A and B, which we apply in mono-therapy at concentrations *a* and *b*, respectively. The concentrations are chosen in a way that the effects are the same (i.e. *E*_*A*_(*a*) = *E*_*B*_(*b*)). All potential combinations of drug A and B (with concentrations (*p · a*, (1 *− p*) *· b*) *∀ p ∈* (0, 1)) could either result in the exact same effect as in mono-therapy (i.e. *E*_*A*_(*a*) = *E*_*B*_(*b*) = *E*_*A,B*_(*p · a*, (1 *− p*) *· b*)) or in a larger (or smaller) effect. He illustrates this with the example of adding 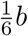 to the dose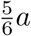. Adding the second drug could, on the one hand, behave as if we would add just another fraction 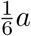 of drug 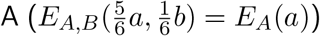, hence adding simply the doses of the two drugs together. On the other hand, adding 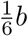 could result in an effect addition with 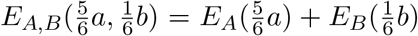, which can be different from *E*_*A*_(*a*), depending on the dose-response relationships of the drugs (see figure 8 in Frei (1913) for the visualisation of this example). He calls the additivity of doses ‘Iso-Additivität’ (iso-additivity) and the additivity of effects ‘Hetero-Additivität’ (hetero-additivity), which would be the same, for a linear relationship of the dose-response curves. He comments that iso-additivity is only possible for drugs with the same effect curves, but he does not mention any assumptions specifically for the target sites of the drugs.

Siegfried Walter Loewe mentioned the concept of additivity first in his work from 1926 (Loewe & Muischnek, 1926), explaining that two drugs would have what he calls a not-varied joint action if they behaved as if two doses of the same drug were combined, which would be ‘additive’. He further explains that this can only be observed for two drugs with the exact same site of action. He uses an isobole-diagramm to explain the concept of additivity (see figure 1 in Loewe and Muischnek (1926) - additivity is given by line number II). An isobole-diagram can be described as follows: consider a graph in which the x-axis displays the concentration of drug A and the y-axis of drug B. Both drugs lead in mono-therapy to a certain effect *e* (i.e. *E*_*A*_(*a*) = *E*_*B*_(*b*) = *e*). An isobole is a line which connects the points *a* and *b*, displaying all the points (all pairs of concentrations 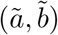) that result in combination in the effect *e*. Two drugs are additive if this line is linear, hence if it includes all points 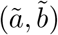) for which holds

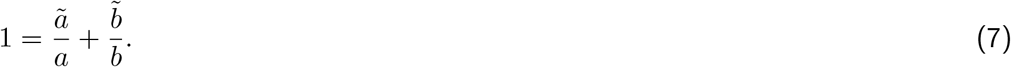

**Figure 1.**
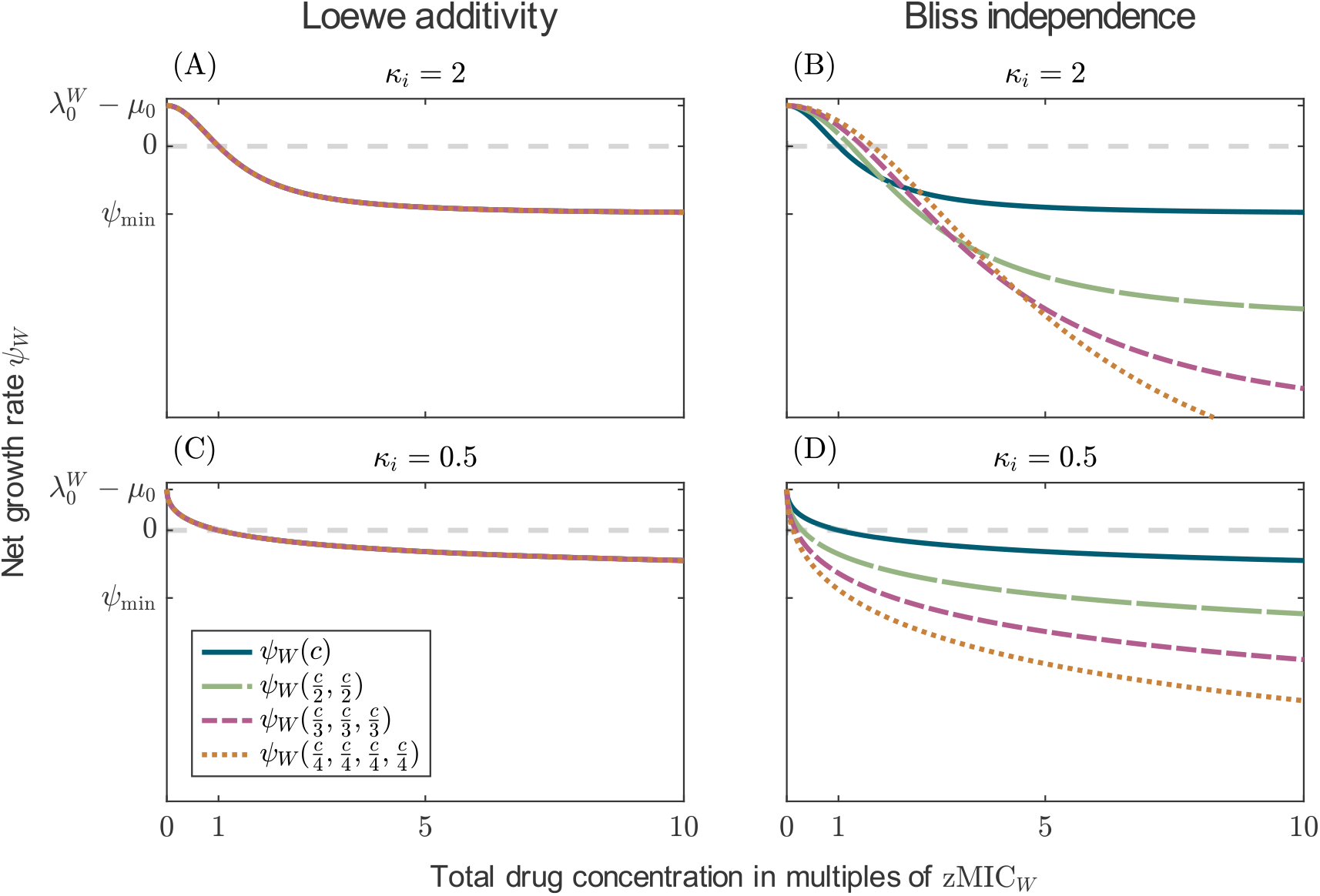
The net growth rate of susceptible bacteria for mono-therapy and combinations with up to four drugs. The functions are displayed for the different additivity models (columns) and different Hill coefficients (rows). For Loewe additivity, all lines coincide. The parameter values are given in table 3, except *ψ*_min_ was set to *−*1*h*^*−*1^.

Equation (7) defines the isobole-equation, which Loewe states two years later in Loewe (1928). Basically, under additivity, a portion of one drug can be replaced by a portion of another drug, without changing the effect, which is essentially ‘iso-additivity’ as described in Frei (1913). Having gotten aware of Frei’s work, Loewe discusses both iso- and hetero-additivity in his work from 1928 and acknowledges that additivity can have multiple meanings: hetero-additivity would not result in a linear isobole and might in fact, look like an antagonism or synergism, which needs to be taken into account when investigating drug-drug interactions for specific combinations.

**Table 2:**
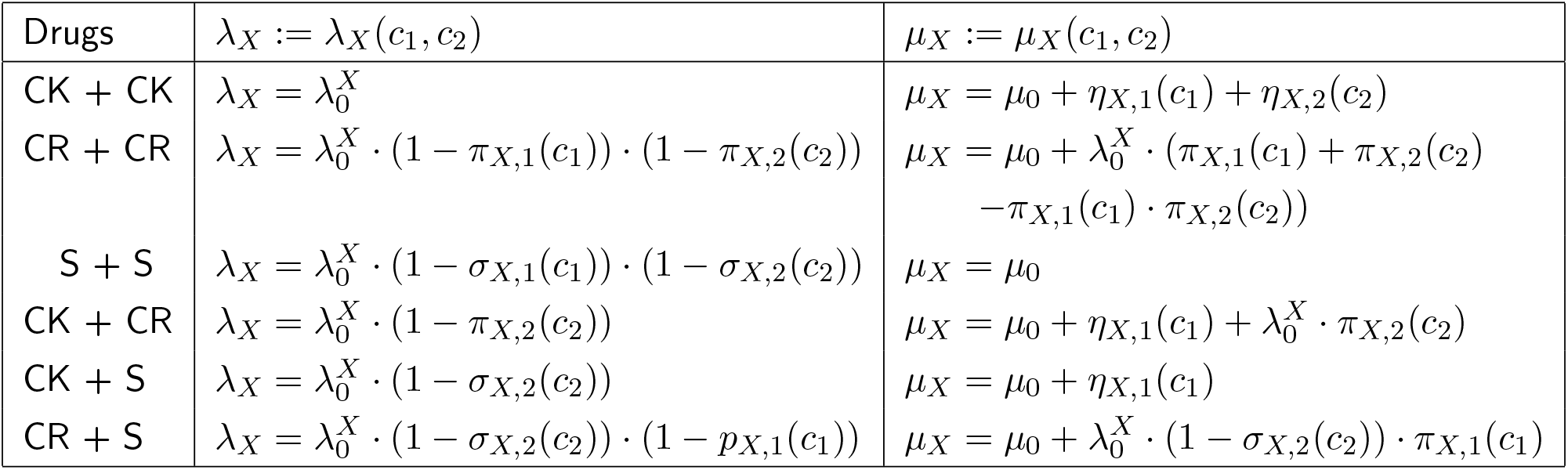
Replication and death rates of a cell with type *X* for the combinations of two drugs with different modes of action assuming Bliss independence. The functions *η*_*X,i*_, *σ*_*X,i*_, and *π*_*X,i*_ with *i ∈* 1, 2 are derived from the action of drug *i* in monotherapy as given in table 1. The parameters describing the drug effect are specific to each drug (i.e. *κ*_*i*_, *ψ*_min,*i*_).

| Drugs | $\lambda_X := \lambda_X(c_1, c_2)$ | $\mu_X := \mu_X(c_1, c_2)$ |
| --- | --- | --- |
| CK + CK | $\lambda_X = \lambda_0^X$ | $\mu_X = \mu_0 + \eta_{X,1}(c_1) + \eta_{X,2}(c_2)$ |
| CR + CR | $\lambda_X = \lambda_0^X \cdot (1 - \pi_{X,1}(c_1)) \cdot (1 - \pi_{X,2}(c_2))$ | $\mu_X = \mu_0 + \lambda_0^X \cdot (\pi_{X,1}(c_1) + \pi_{X,2}(c_2) - \pi_{X,1}(c_1) \cdot \pi_{X,2}(c_2))$ |
| S + S | $\lambda_X = \lambda_0^X \cdot (1 - \sigma_{X,1}(c_1)) \cdot (1 - \sigma_{X,2}(c_2))$ | $\mu_X = \mu_0$ |
| CK + CR | $\lambda_X = \lambda_0^X \cdot (1 - \pi_{X,2}(c_2))$ | $\mu_X = \mu_0 + \eta_{X,1}(c_1) + \lambda_0^X \cdot \pi_{X,2}(c_2)$ |
| CK + S | $\lambda_X = \lambda_0^X \cdot (1 - \sigma_{X,2}(c_2))$ | $\mu_X = \mu_0 + \eta_{X,1}(c_1)$ |
| CR + S | $\lambda_X = \lambda_0^X \cdot (1 - \sigma_{X,2}(c_2)) \cdot (1 - p_{X,1}(c_1))$ | $\mu_X = \mu_0 + \lambda_0^X \cdot (1 - \sigma_{X,2}(c_2)) \cdot \pi_{X,1}(c_1)$ |

**Table 3:** Parameter descriptions and their values used to generate the results. When multiple values are given, these were varied during the analysis. The pharmacodynamic parameter values were chosen in accordance to values measured from experimental data (Regoes et al., 2004). Values for the cost and benefits of mutations were reviewed by Igler et al. (2021); the values chosen here specifically rely on the work by Melnyk et al. (2015) and Spohn et al. (2019). We consider a wide range of mutation probabilities, including values on the extreme high and extreme low end. Examples of these values can be found in Rodríguez-Rojas et al. (2014), Imhof and Schlötterer (2001), Oliver et al. (2004), McGrath et al. (2014), and Wang et al. (2001). Please note that we use the same zMIC_*W*_ for all drugs. However, as the concentration range is always displayed in multiples of the zMIC_*W*_, the results are independent of the actual values, which could differ between drugs.

| Parameter | Description | Value |
| --- | --- | --- |
| $N_0$ | initial population size | $10^{10}$ |
| $\lambda_0$ | intrinsic replication rate | $0.7h^{-1}$ |
| $\mu_0$ | intrinsic death rate | $0.1h^{-1}$ |
| $u_i$ | mutation probability, resistance to drug i | $10^{-11}, 10^{-9}$ or $10^{-5}$ |
| $zMIC_W$ | wild-type MIC | $1 \frac{\mu g}{ml}$ |
| $\beta_{res}$ | benefit of mutation in terms of resistance | 28 |
| $\gamma_i$ | cost per mutation | 0.1 |
| $\psi_{min}$ | growth rate at very high concentrations: | |
| | $\psi_{min}^{CK}$ | $-6h^{-1}$ |
| | $\psi_{min}^{CR}$ | $-0.73h^{-1}$ |
| | $\psi_{min}^S$ | $-0.065h^{-1}$ |
| $\kappa$ | Hill coefficient | 0.5, 1, or 2 |

Chester Ittner Bliss defines ‘independence’ (he calls it ‘independent joint action’) through stochastic independence, specifically referring to drugs with different toxic actions (Bliss, 1939). The proportion of cells surviving the treatment with two drugs (drug A and B) is given by the product of the proportions surviving the treatment with either one of the drugs:

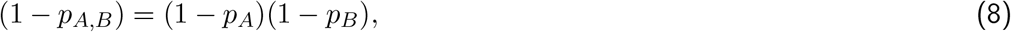

where *p*_*A*_ and *p*_*B*_ are the proportions of cells killed by the treatment with either drug A or drug B, respectively, and *p*_*A,B*_ is the proportion not surviving the combined treatment. Hence, the two drugs act independently of each other. He additionally defines the concept of ‘similar joint action’, which is essentially (iso-)additivity, as described by Frei and Loewe, for two drugs targeting the same set of receptors. As Bliss independence is defined on the surviving fraction of a population, not on the net growth rate of cells, recent modelling studies often refer to the definition of Bliss independence on rates as given by Baeder et al. (2016). In this definition, the joint kill rate of the combination is given by the sum of the individual kill rates of the drugs, which results in an effect-addition, similar to the description by Frei under the name hetero-additivity.

Both, Bliss and Loewe make a clear assumption on the target sites of the drugs for which their models apply. Nevertheless, the models are often seen as rival approaches in which one should universally be favoured over the other. Greco et al. (1995), for example, use the so-called ‘sham experiment’ to argue for Loewe additivity and against Bliss independence. The experiment is described like this: one aims to measure the joint effect of two drugs, but when adding the dose of the second drug one accidentally takes another dose from the first drug. The results of the measurement would show what ‘no interaction’ would look like. The additivity model should accurately capture this and predict no interaction when combining a drug with itself, which is fulfilled by Loewe additivity. Bliss independence is criticised for not fulfilling the sham experiment. Independent of the underlying assumptions on the modes of action, the authors therefore generally favour Loewe additivity over Bliss independence.

However, this brief excursion into the literature shows that neither of the models is universally applicable and rather complements the other than opposes it. Both models were defined for specific assumptions on the modes of action. This is further supported by the work of Baeder et al. (2016). The authors simulated the combined drug effect of two drugs with a multi-hit model, explicitly simulating the drug-target binding. Bliss independence (defined on rates) was able to predict the simulated drug effect for drugs with different target sites and Loewe additivity for drugs with the exact same target site.

#### 2.3.2 The implementation of Bliss independence and Loewe additivity

Following the assumption for the sites of action of the drugs, Bliss independence can be implemented for all combinations of drugs differing in their dose-response curves or modes of action. For the joint effect of the drugs on the kill rate, we follow Baeder et al. (2016) who showed that Bliss independence translates into additivity of kill rates. For the effect on the replication rate, we assume stochastic independence, i.e. probabilities for successful replication multiply. Table 2 shows the resulting replication and death rates for combining two drugs with the same or different modes of action. We would like to point out that neither of these implementations leads to stochastic independence of survival probabilities (based on our derivation of extinction probabilities in appendix C) and thus does not actually match the original definition by Bliss (1939); see appendix B.

For Loewe additivity, since drugs act on the same target site, we consider only combinations of drugs with the same mode of action. To calculate the joint drug effect, we follow the implementation of Lederer et al. (2018). For two drugs which have in mono-therapy the net growth rate functions *ψ*_*X*,1_(*c*_1_) and *ψ*_*X*,2_(*c*_2_) for a type *X*, the net growth rate under the combination treatment can be calculated by

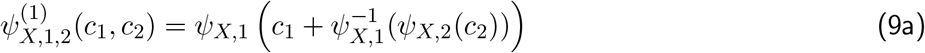

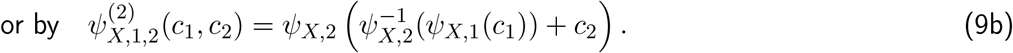

Equations (9a) and (9b) lead to the same results only under the condition that the dose-response curves are the same (up to potentially different minimal inhibitory concentrations), which we assume in the following when considering Loewe additivity.

To compare different treatments (mono-therapies and combinations), we scale the individual drug concentrations with the respective MICs and effectively keep the total drug concentration constant. More precisely, we choose the individual concentrations *c*_*i*_ such that 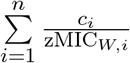 is the same for all treatments, irrespective of the number and types of drugs in the treatment. Drug ratios refer to the scaled concentrations. E.g., for a two-drug combination with a 1:1 ratio, we have *c*_1_*/*zMIC_*W*,1_ = *c*_2_*/*zMIC_*W*,2_. For notational simplicity and concreteness, we set 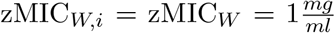 for all drugs in the following, but results apply more generally.

Under Loewe additivity, the net growth rate is independent of the number of drugs (displayed for CK drugs in figures 1A and C). For Bliss independence (panels B and D), by contrast, the number of drugs matters. Two observations can be made. First, the total concentration at which the net growth rate is zero (the MIC of the combination) depends on the number of drugs and the maximum strength of the drug effect *ψ*_min_ (figures 1A and C, and S1, S2). Interestingly, the increase in the concentration required for growth inhibition, which we see for drugs with *κ* = 2 and low *ψ*_min_ (i.e. |*ψ*_min_| large), can be observed in experiments (Russ & Kishony, 2018). Unfortunately, the study did not consider drugs with shallow dose-response curves. Therefore, we do not know if the decrease in the concentration needed for growth inhibition that we observe for those curves can be observed empirically as well.

The second observation from figure 1 is that the growth rate in the limit of high concentrations decreases with an increasing number of drugs since we are summing up the kill rates of the drugs. It seems rather unlikely that the effect of multiple drugs is so much stronger than that of a single drug in isolation and that this effect increases indefinitely with the number of drugs, which shows that there are puzzles with translating Bliss independence to rates. We come back to this in the discussion section. For combinations of drugs with the other modes of action, the maximum effect can also increase with the number of drugs, but the minimum growth rate of susceptible bacteria is limited to *−μ*_0_ for bacteriostatic drugs and *−μ*_0_ *− λ*_0_ for bactericidal drugs that act during replication.

Examples of dose-response curves (net growth rates) for the resistant cell types and for different drug combinations are shown in figures S3-S8.

### 2.4 The probability of treatment success

For the comparison of different treatments, we calculate the probability of treatment success, which we define as the extinction probability of a bacterial population of initially *N*_0_ cells undergoing treatment.

#### 2.4.1 Exact calculation of the probability of treatment success

In our model, cells replicate and die independently from each other (see section 2.1). Hence, each cell in the initial population founds a lineage with its own independent fate – extinction or growth. To calculate the extinction probability of the entire population, we need to multiply the extinction probabilities of all these independent lineages. For the treatment with *n* drugs, the probability of treatment success is thus given by

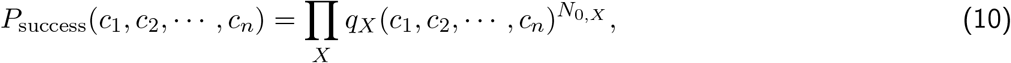

where *q*_*X*_(*c*_1_, *c*_2_, *· · ·, c*_*n*_) is the extinction probability of a cell lineage starting with a cell of type *X* (including the possibility of *de novo* mutations) and *N*_0,*X*_ cells of type *X* are present in the initial population. Importantly, even cells with a positive net growth rate might fail to establish a long-term lineage due to stochasticity at low cell numbers. The extinction probabilities *q*_*X*_ are derived in appendix C.

#### 2.4.2 Approximation of the probability of treatment success

Equation (10) allows us to compare the efficiencies of different treatments but we cannot see from it why a certain treatment is better than another. We therefore derive an approximation with which we can disentangle where the benefit of a treatment comes from, using an approach that is common in models of evolutionary rescue (see e.g. Bell & Collins, 2008; Orr & Unckless, 2008; Martin et al., 2013; Uecker et al., 2014; Czuppon et al., 2021). In the main text, we only give the derivation for mono-therapy; the extension to combination treatment with two drugs is given in appendix D.

We first focus on the *de novo* evolution of resistance. Two factors determine whether the population adapts or goes extinct: the number of resistant mutants appearing by mutation during treatment and the probability that these mutants establish, i.e. do not suffer stochastic loss. New mutants appear at rate *u · λ*_*W*_ (*c*) *· W* (*t*), where *W* (*t*) is the wild-type population size at time *t*. Given the large initial population size, we approximate the decline of the wild-type deterministically:

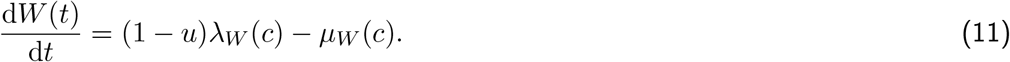

Since the appearance of mutants follows a Poisson process, the number of mutants over the time course of the whole treatment is given by a Poisson distribution with mean

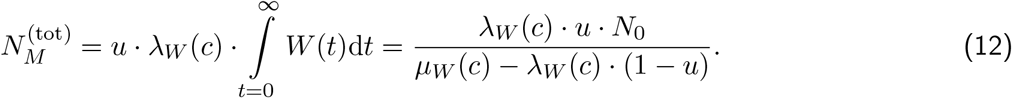

Each of them establishes a long-term lineage with probability

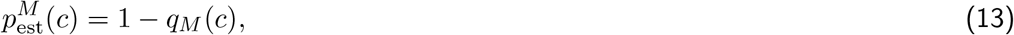

with *q*_*M*_ being the extinction probability as derived in appendix C. The probability of treatment success, i.e. the probability that no successful mutant appears over the time course of the treatment, is then given by the zeroth-term of a Poisson distribution with mean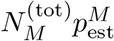:

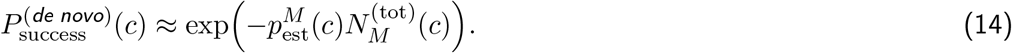

We hence see that the probability of treatment success is determined by the product of the total number of new mutations appearing and their establishment probability. Single mutants in the standing genetic variation are eradicated with probability 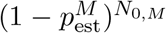. A treatment works well if both 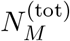 and 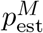 are small, where the importance of minimising the establishment probability increases with the number of pre-existing mutants. By calculating these terms and comparing them among treatments, we can gain insights into whether a treatment is particularly successful because it prevents the appearance of new mutants or minimises the establishment probability or both.

## 3 Results

To identify the optimal conditions for antibiotic combinations, we will compare in this section the success probability of different combination treatments, including a comparison to monotherapy at the same total dose. We evaluate the treatment efficiency for several mutation probabilities towards resistance, ranging from low to high, and in the absence and presence of pre-existing mutants. We do so across a large range of concentrations. Absence of pre-existing mutations is unlikely, unless mutation probabilities are very low. Yet, it is instructive to consider this scenario, and it can be interpreted as preventing the *de novo* evolution of resistance. The parameter values used for the numerical analysis are displayed in table 3. Examples of drugs and drug combinations that are used to treat infections, together with key properties relevant to out study, are listed in table S1. Whenever possible, we derive results for both additivity models (Bliss and Loewe) to evaluate if they lead to the same conclusion. We first explore the role of the mode of action on the evolution of resistance in mono-therapy. We then compare the efficiency of mono-therapies with combinations and subsequently focus on combinations of drugs with different modes of action. Lastly, we extend the comparison to treatments at which the drugs are administered at different ratios and consider combinations with more than two drugs.

### 3.1 Comparing mono-therapies of drugs with different modes of action

Before considering multiple drugs in combination, we first compare drugs with different modes of action in mono-therapy.

#### Bacteriostatic drugs limit the de novo evolution of resistance better than bactericidal drugs if both have the same dose-response curve

First, we assume that the drugs have the same effect on the net growth rate, shown in figure 2A. Panel B displays the probability of treatment success for a fully susceptible population and a population with one pre-existing mutant. Let us focus for now on the evolution of resistance in the susceptible population. Panel B (left) reveals that the bacteriostatic drug leads to a higher probability of treatment success than the bactericidal drugs and that the bactericidal drug acting during replication (CR) is better than a bactericidal drug which kills independent of replication (CK). To explain the difference in treatment efficiency, we compare the effect of the treatments on the number of mutants arising *de novo* and their establishment probability. Figure 2C shows that fewer mutants are generated during the treatment with the bacteriostatic drug compared to the bactericidal drugs (S *<* CR *<* CK). However, bactericidal drugs are better at preventing mutants from establishing, with the order being reversed, i.e. CK *<* CR *<* S. A mathematical proof shows that these observations generalize beyond the numerical example (appendix E): The more drug effect is allocated on reducing the replication rate vs increasing the death rate, the lower the number of new mutants, the higher the establishment probability, and the higher the total probability of treatment success. I.e., the effect on the number of mutants outweighs the effect on the establishment probability.

**Figure 2.**
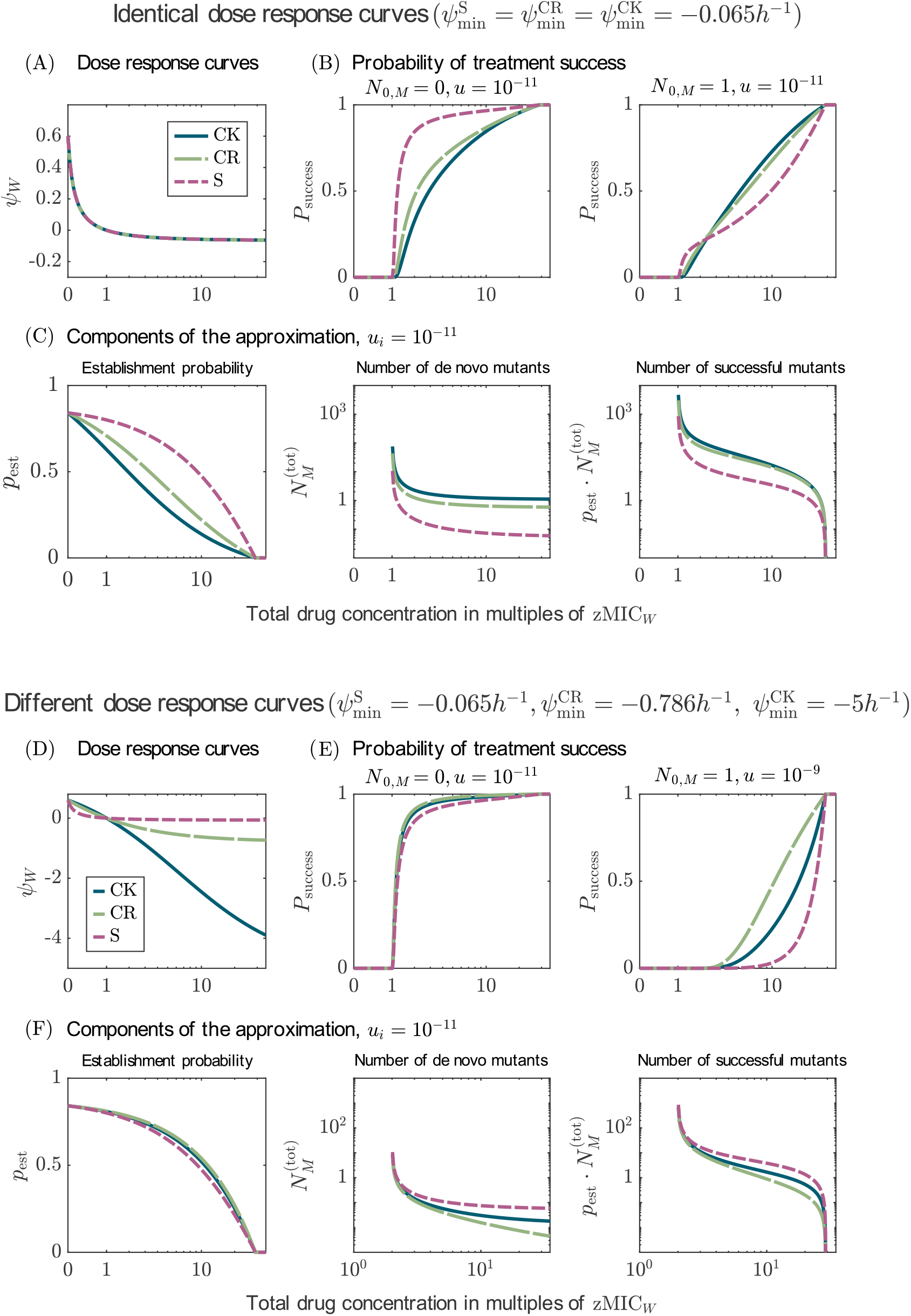
Comparison of mono-therapies with drugs with different modes of action. Panels A-C show the results for a comparison of drugs with the same dose-response curve (*ψ*_min_ = *−*0.065*h*^*−*1^), and panels D-F show the the results for drugs with different dose-response curves (differing in *ψ*_min_). Panels A and D display the respective dose-response curves. Panels B compare the probabilities of treatment success for the three drugs in the absence and presence of pre-existing mutants. Panels E show the probabilities of treatment success for different mutation probabilities. Panels C and F show the terms used in the approximation of the probability of treatment success (equation (13) and (12) and their product). The results are shown for *κ* = 1 and *u* = 10^*−*11^ (except for panel E (right)).

#### Bactericidal drugs are better than bacteriostatic drugs at preventing mutants from establishing, which can turn them into the better option when mutants pre-exist prior to treatment

When mutants pre-exist in the population before the treatment, the effect on the establishment probability gains in importance. Figure 2B (right) shows that for drug concentrations above ca. 2.5 *·* zMIC_*W*_, the bactericidal drug acting independent of replication leads to the highest probability of treatment success and the bacteriostatic drug to the lowest. At such high concentrations, *de novo* mutants are rare anyway, making it more important to prevent pre-existing mutants from establishing than to further reduce the number of new mutations. Note that establishment probabilities are generally high for the chosen parameter values such that even a single pre-existing mutant has a strong visible effect. For low concentrations, the number of new mutants is high (panel C), and efficiently decreasing the number of mutants by using a bacteriostatic drug is still the better strategy.

#### Bactericidal drugs can become better than bacteriostatic drugs for high enough bactericidal effects even in preventing the de novo evolution of resistance

In reality, the value of *ψ*_min_ of the bactericidal drugs is most likely lower than that of the bacteriostatic drug; this especially holds for the CK drug. We therefore include such a drug comparison (figure 2D). In this comparison, the bacteriostatic drug becomes the worst option since the other two drugs lead to fewer new mutants (panels F). At first sight, surprisingly, the mutant establishment probability is higher now for the bactericidal drugs than it was for the high value of *ψ*_min_ (i.e. |*ψ*_min_| small). This can be understood by considering the dose-response curves: below the MIC, bactericidal drugs with a lower *ψ*_min_ have a *higher* net growth rate (panels A and D show this for the wild type, but the same applies to the mutant). Overall, using a drug with a mode of action that strongly decreases the number of mutants, even at the cost of allowing resistant mutants to establish more easily than with other drugs, seems to be a good treatment strategy when resistance evolves *de novo*. When mutants are present in the initial population, using a drug that reduces the mutants’ establishment probability most efficiently is better.

### 3.2 Comparing mono-therapies with two-drug combinations at equal drug ratios

We compare in this section the treatment efficiency of mono-therapy with that of combination treatment with two drugs, given a constant total dose. For combination therapy, the total drug dose is thereby split equally between the two drugs (for unequal MICs, the drug doses are scaled with the respective zMIC_*W*_, see model description). We will, for now, consider combinations in which all drugs of the comparison have the same dose-response curves and focus on the influence of the Hill coefficient *κ*. We mainly present results for bactericidal drugs and only show specific cases for drugs with different modes of action since the results are otherwise the same. All results are shown for both models of drug additivity (Loewe and Bliss). We also varied the intrinsic replication and death rate and *ψ*_min_. The results are not shown, as the change in parameter values did not affect the outcome of the comparison.

#### Combination therapy is always better than mono-therapy for drugs with shallow dose-response curves

Figure 3A-C shows that the probability of treatment success is always higher for combination treatment than for mono-therapy when the dose-response curves are shallow (i.e. *κ*_*i*_ *≤* 1) (the results for *κ*_*i*_ = 1 are shown in figure S9). This result is independent of the mutation probability and the pre-existence of mutants prior to treatment (figure S10A-F). It holds both for combinations displaying Bliss independence and combinations displaying Loewe additivity. Combinations with Bliss independence lead to a higher (or equal) probability of treatment success than drugs with Loewe additivity. These observations are the same for all modes of action of the drugs (see figures S11A-F and S12A-F).

**Figure 3.**
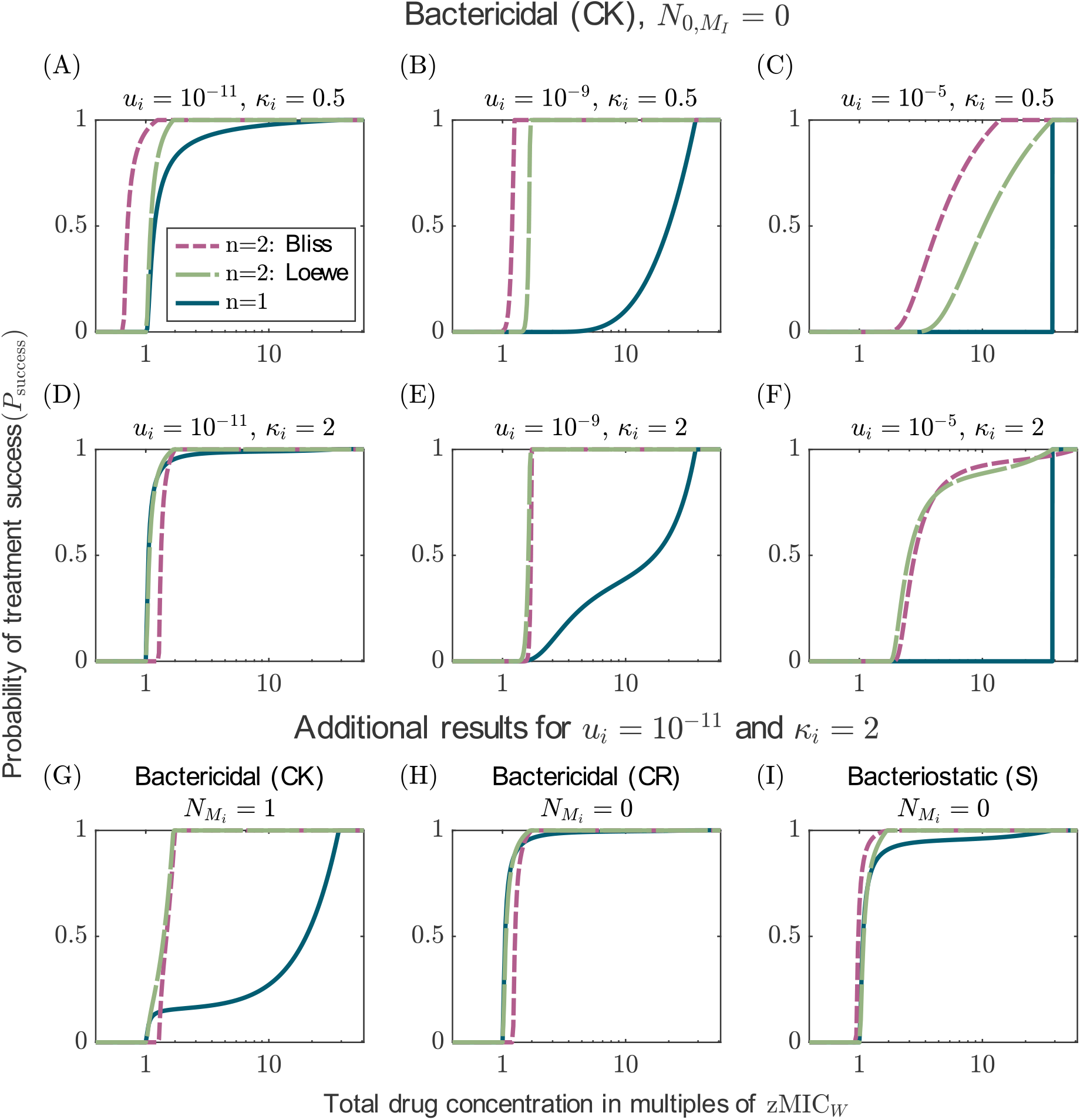
Comparison of the probabilities of treatment success under mono-therapy (*n* = 1) and a two-drug combination (*n* = 2). The results of the two-drug combination are shown for both additivity models, Bliss independence (pink) and Loewe additivity (green). Panels A-F display the results for bactericidal drugs acting independent of replication. Each column displays the comparison for different mutation probabilities (see panel header of the first row) and each row for different Hill coefficients (see values of *κ* in the header). Panels G-I display additional results for drugs with *κ*_*i*_ = 2 and *u*_*i*_ = 10^*−*11^ (i.e. the parameter settings of panel D). Panel G considers the pre-existence of one single mutant per type in the initial population and panel H and I display the results for the other two modes of action.

What is causing the advantage of combination treatment over mono-therapy? Dissecting the problem following the rationale of approximation (14) shows that for the chosen parameters, the number of new single mutants (for combination therapy, the sum of both single mutants) that appear during treatment are similar for both treatments, yet their establishment probabilities are much lower under combination therapy (see figure S13). The establishment probability of the double mutants is similar to that of the single mutant in mono-therapy but they are rarely generated. One can also understand this by considering the dose-response curves of all types (figure S3): both susceptible cells and single mutants have lower net growth rates under combination therapy than under mono-therapy. For *κ ≤* 1, double mutants have similar or lower growth rates under combination therapy than under mono-therapy (but as said before, they are rare). Combination therapy is thus better than mono-therapy by decreasing the establishment probability of the single mutants and increasing the genetic barrier to resistance. The lower establishment probability of the single mutants is also intuitively clear beyond the numerical example: in mono-therapy, they effectively experience a concentration that is lower by a factor *β* (which we set to 28, see table 3) than the one experienced by the wild type. In combination therapy, they still experience 1/2 of the second drug to which they are susceptible.

#### Mono-therapy can sometimes be better than two-drug treatments for drugs with steep dose-response curves

For a combination of drugs with steep dose-response curves, we can sometimes observe small concentration ranges in which mono-therapy leads to a higher probability of treatment success than combination therapy (figure 3D-I). First, monotherapy can be slightly better than drug combinations displaying Bliss independence for concentrations slightly above the MIC (zMIC_W_) of the wild-type (panels D,G,H). This can be explained by the increased MIC for combinations that we already saw in figure 1 (see also figure S3) – slightly above zMIC_W_, mono-therapy can eradicate the wild-type while combination therapy cannot. As briefly indicated above, whether this shift in MIC occurs or not, depends on the *ψ*_min_ of the drugs. For the parameter values used here, the shift in MIC occurs – opening up a small window in which mono-therapy is superior – for both bactericidal drugs, but not for the bacteriostatic drugs (panels D,H,I; see figures S3, S4, and S5).

Second, under Bliss independence, mono-therapy leads to higher treatment success than combination therapy when drug concentrations are around *β*_res_ *·* zMIC_W_ (the MIC of single mutants in mono-therapy) and mutation probabilities are high. In this regime, concentrations in mono-therapy are so high that even resistant types are controlled well. In contrast to drugs with shallow dose-response curves, double resistant types in combination therapy have a higher growth rate than single mutants in mono-therapy at those concentrations and can still grow (see figure S3 for the growth rates of all types). At the same time, they arise at a substantial frequency due to the high mutation probability. This effect is either absent or negligibly small for the other modes of action (figures S11, S12).

When mutants pre-exist, mono-therapy is usually much less effective, which further reduces the drug ranges in which mono-therapy is better than combination treatment (see figure 3G and figure S10 for more results). An important exception to this are scenarios where mutation probabilities are high (which also implies many pre-existing mutants). In this case, monotherapy is superior to two-drug combinations with Bliss independence because types that are fully resistant to the respective treatment are better controlled by mono-therapy than by combination therapy (figure S10).

To conclude, we see that although there are some noteworthy exceptions, the two-drug combination seems to be overall a better treatment choice than mono-therapy across different mutation probabilities and for all three modes of action, despite of the decrease in the individual drug doses.

### 3.3 Comparison of combinations of two drugs with different modes of action (50:50 ratio)

In this section, we discuss combinations of drugs which differ in their modes of action. For the comparison, we will again match the dose-response curves of the individual drugs as in section 3.1. The dose-response curves of the combination (i.e. the function *ψ*(*c*_1_, *c*_2_)), however, differ between the modes of action of the drugs (figure S7). As in the previous section, the total drug concentration is split equally between the two drugs. We will first compare combinations of drugs with different modes of action to combinations of drugs with the same mode of action to evaluate if mixing drugs of different modes of action is generally beneficial. Then, we compare various combinations of drugs with different modes of action to identify which one is optimal. We show results for drugs with a Hill coefficient *κ*_*i*_ = 1. Additional results for other parameter choices can be found in SI section S4. Varying the intrinsic replication and death rate and the Hill coefficients does not change the outcome of the comparisons.

#### Combining drugs with different modes of action can sometimes be the best option by reducing both the number of mutants and the establishment probability

We find that using drugs with different modes of action can lead to a higher probability of treatment success than treatments using two drugs with the same mode of action. This is shown in figure 4A-C: combining a bacteriostatic drug with a bactericidal drug acting independent of replication (S+CK) is better than using two bacteriostatic drugs (S+S) and at least as good as combining two CK drugs. To see what makes this combination so successful, we can look at the number of mutants generated and their establishment probabilities; this is shown in panels D-F for a high mutation probability where the CK+S combination is better than both of the other combinations. Combining two bactericidal drugs leads to the lowest establishment probability but the highest number of mutants, and combining two bacteriostatic drugs leads to the lowest number of mutants but the highest establishment probability (panels D and E). By suppressing both factors fairly well, the mixed treatment becomes the best treatment option. Hence, in this particular case, combining different modes of action is advantageous. However, a similar comparison with a bacteriostatic (S) and a bactericidal drug acting during replication (CR) shows that the mixed treatment (S+CR) never results in the highest probability of treatment success (figure S14).

**Figure 4.**
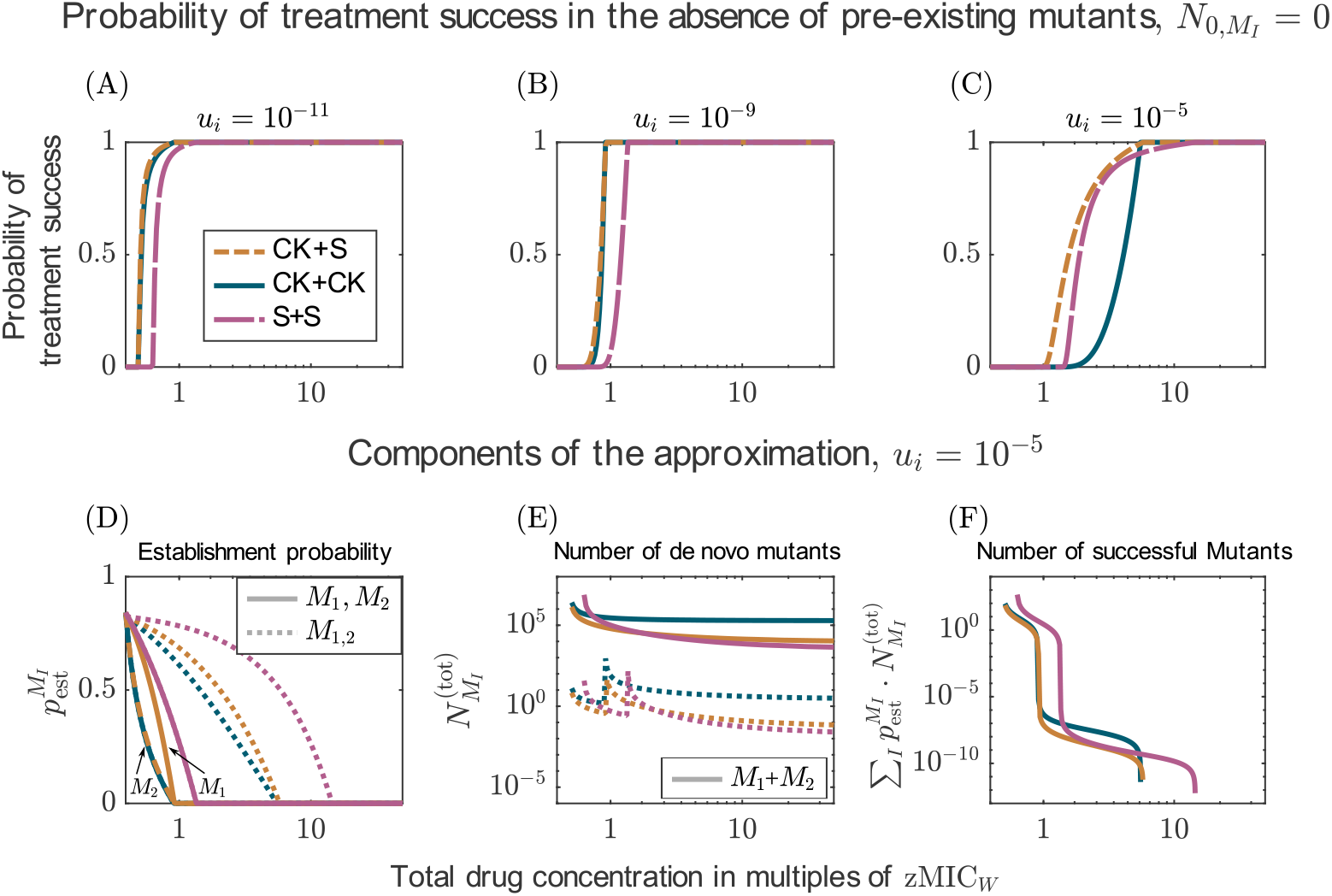
Comparison of the probabilities of treatment success for combinations of drugs with either the same (S+S and CK+CK) or different modes of action (S+CK). The upper row shows the probabilities of treatment success for three different mutation probabilities. The lower row shows the components of the approximations of the probability of treatment success (equations (14), (A16), and (A21)) for the highest mutation probability (*u*_*i*_ = 10^*−*5^). All drugs have the same dose-response curve with Hill coefficient is *κ*_*i*_ = 1 and *ψ*_min_ = *−*0.065*h*^*−*1^.

#### Combining a bacteriostatic drug with a bactericidal drug can be both the best or worst treatment decision

Figure 5 compares combinations of two drugs with different modes of action. The best treatment option, in the absence of pre-existent mutants, is for all mutation probabilities the combination of the bactericidal drug acting independent of replication (CK) with a bacteriostatic drug (S) (figure 5A-C). When mutants pre-exist in the initial population, combining drugs with the two bactericidal modes of action (CK+CR) is sometimes slightly better (panels D-F). Combining the bactericidal drug acting during replication (CR) and the bacteriostatic drug (S) is always the least good option. Results are similar for drugs with Hill coefficients *κ*_*i*_ = 0.5 and *κ*_*i*_ = 2 with the noteworthy exception that the S+CR combination can be better than the CR+CK combination if *κ*_*i*_ = 2 and the mutation probability high (figures S15 and S16). All combination treatments, even the worst one, are better than the best mono-therapy (see grey line in the plots).

**Figure 5.**
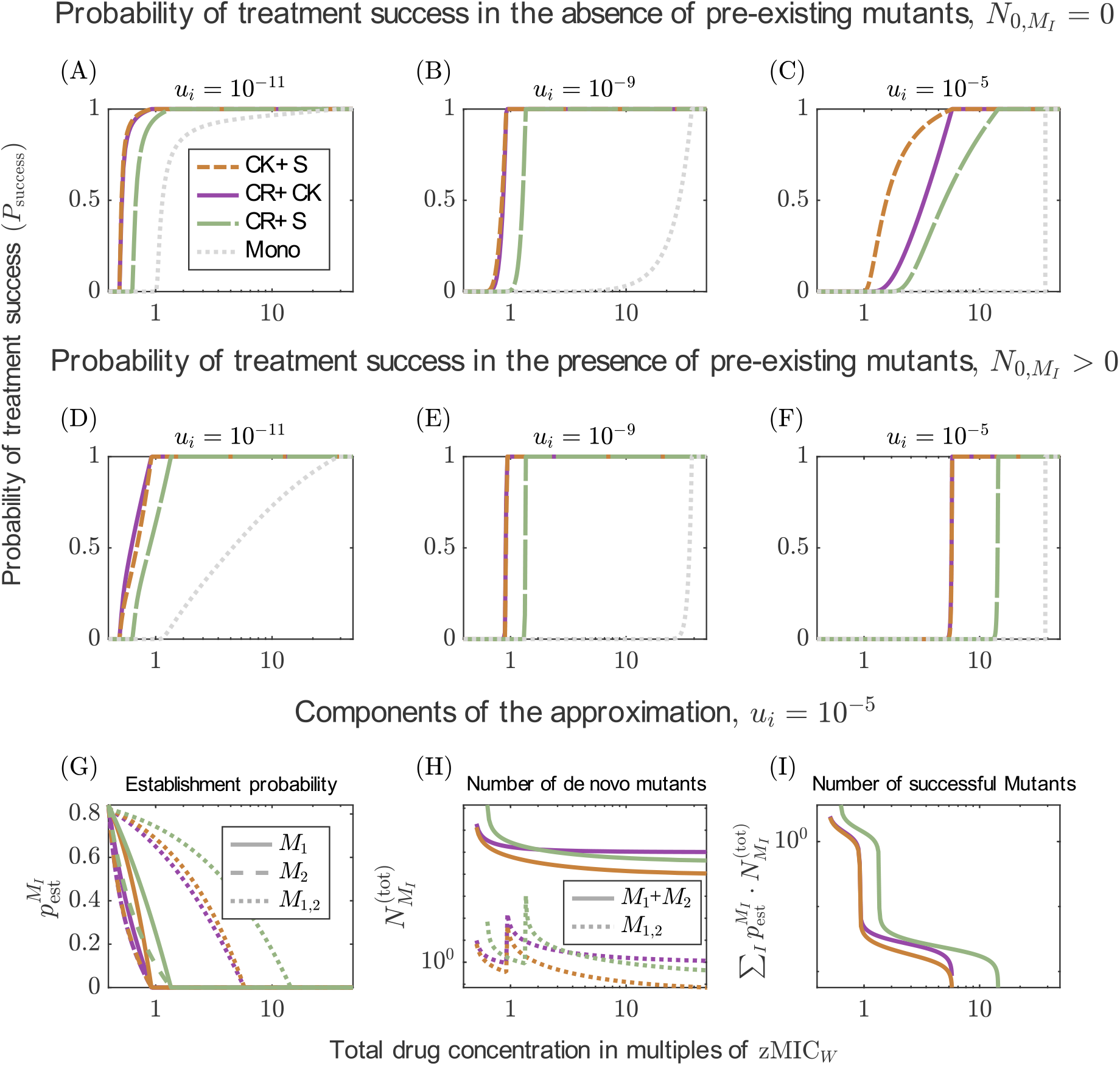
Comparison of the probabilities of treatment success for various combinations of drugs with different modes of action and for the best mono-therapy. The first and second row show the probabilities of treatment success for three different mutation probabilities in the absence (A-C) and in the presence (D-F) of pre-existing mutations. As reference, the best mono-therapy is displayed in grey. The third row shows the components of the approximations of the probability of treatment success (equations. (14), (A16), and (A21)) for the highest mutation probability (*u*_*i*_ = 10^*−*5^). All drugs have the same dose-response curve with Hill coefficient is *κ*_*i*_ = 1 and *ψ*_min_ = *−*0.065*h*^*−*1^.

We can again look at the two factors that determine resistance evolution – appearance and establishment of mutants (panels G-I). Combining the bacteriostatic drug with the bactericidal drug acting independent of replication (CK+S) leads to the lowest number of mutants. This combination suppresses the establishment probabilities as much or almost as much as the combination of two bactericidal drugs, which has the lowest establishment probabilities. For the combination of the bacteriostatic drug with the bactericidal drug acting during replication (S+CR) is bad in terms of establishment probabilities and – somewhat surprisingly – also in terms of number of mutants generated. The result does not change much when the first drug (the drug that is first written in the legend) has a higher maximum effect, i.e. lower *ψ*_min_ (figure S17).

Overall, combining drugs with different modes of action can contain both the establishment probability of mutants and the number of mutants, resulting in a more efficient treatment than a combination that mainly influences one of the two factors. However, not all combinations of modes of action are equally good. Combining a bacteriostatic drug with a bactericidal drug killing independently of replication (S+CK) is the best (or very close to the best) treatment option across all comparisons. Combining a bacteriostatic drug with a bactericidal drug that acts during replication (S+CR) is one of the worst combinations. However, it is still much better than the best mono-therapy.

### 3.4 Comparison of two-drug combinations at unequal drug ratios

In the previous sections, we discussed comparisons of treatments in which drugs were either administered in mono-therapy or in a two-drug combination in which the total dose was equally split between the two drugs. In this section, we want to extend the comparison of the previous sections to combinations in which the drugs are not administered at a 50:50 ratio. We will provide an overview of different examples to show how the drug characteristics can influence the outcome of combinations administered at uneven drug ratios. The results in this section are displayed for drugs with Bliss independence; more results can be found in SI section S5.

#### The ideal drug ratio might vary across the concentration gradient and across mutation probabilities even when combining drugs with the same drug characteristics

For the combination of drugs with the same shallow dose-response curves (*κ*_*i*_ *≤* 1), we can see, independent of the total drug dose, the mode of action, the mutation probability, and the pre-existence of resistance mutations, that the two drugs should be given at an equal ratio to maximise the probability of treatment success (see panels A-C in figure 6 and figures S18-S20, for drugs with the other two modes of action and in the presence of preexisting mutants). For drugs with steep dose-response curves, mono-therapy is advantageous at low concentrations for low mutation probabilities (panels D and E) and high concentrations for high mutation probabilities (panel F). Both results – those for shallow and those for steep dose-response curves – align with our comparison of mono-therapies vs two-drug combinations in section 3.2. In addition to the explanations provided there, we show in figure S6 for all types the ranges where net growth rates are positive/negative.

**Figure 6.**
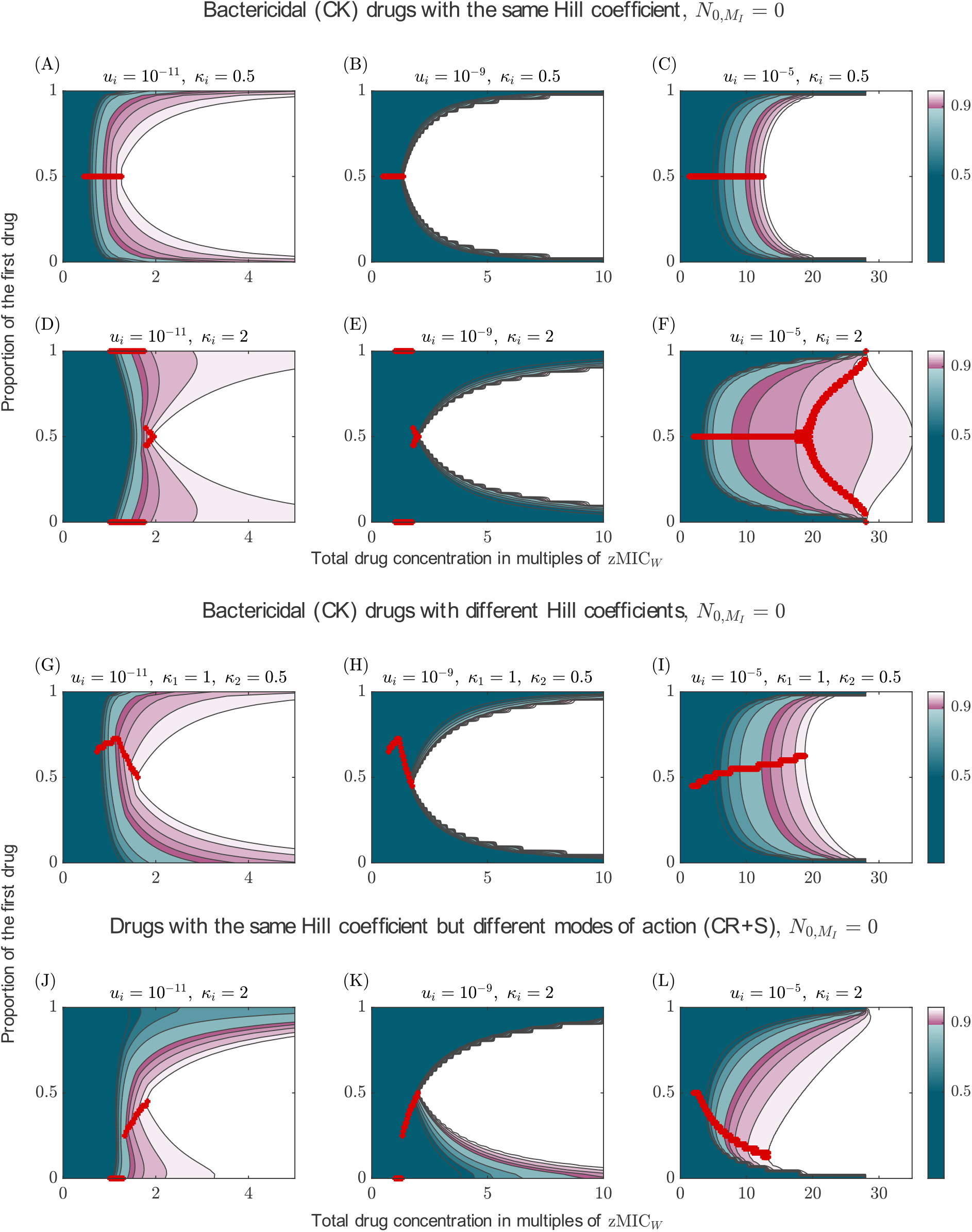
Probability of treatment success for combinations of two drugs administered at unequal ratios. The *x*-axis gives the total drug concentration, the *y*-axis the proportion of the first drug, and the color indicates the probability of treatment success. The white area indicates almost certain treatment success (*P*_success_ *≥* 99%). The red marker indicates the ratios which maximize the probability of treatment success for a specific concentration. The optimal ratio is not shown for cases in which all treatments are equally bad, or once the first treatment has reached a success probability of at least 99%. The figure provides an overview over specific cases (see row titles), more results can be found in SI section S5. Results are shown for different mutation probabilities, as displayed in the column header. Panels A-F show results for combination of drug with the same Hill coefficient (either *κ*_*i*_ = 1 or *κ*_*i*_ = 2. Panels G-I display results for a combination of drugs with different Hill coefficients (*κ*_1_ = 1, *κ*_2_ = 0.5) and panels J-L for drugs with different modes of action (both with *ψ*_min_ = *−*0.1*h*^*−*1^, but the same Hill coefficient *κ*_*i*_ = 2).

We can further ask which drug ratio minimises the concentration needed to have an almost certain treatment success, which we define here as *P*_success_ *≥* 99% (the white area in the panels of figure 6). For shallow dose-response curves, the minimum concentration is lowest for a 50:50 ratio (the boundary between the colored and white areas is concave). For steep dose-response curves, a 50:50 is optimal for low mutation probabilities. In contrast, for high mutation probabilities, mono-therapy minimises the concentration required for almost certain treatment success, and in combination treatment, much higher concentrations would be needed to achieve a treatment success of 99%. Yet, at the concentration where mono-therapy gives a 99% probability of treatment success, administering drugs at a 50:50 ratio still leads to *P*_success_ *≈* 96%.

When mutants pre-exist, choosing the drug ratio that minimises the concentration required for an almost certain treatment success (the boundary between the colored and white areas) becomes more important as the success probability changes more drastically with the drug ratio (figure S20).

For drugs combining according to Loewe additivity, a 50:50 ratio always minimises the concentration required for almost certain treatment success (figure S21).

#### The optimal drug ratio varies when combining drugs with different drug characteristics

When combining drugs with different Hill coefficients (figure 6G-I) or different modes of action (panels J-L), it can be beneficial to administer two drugs at unequal doses; this is especially true if mutation probabilities are high. Unlike for a combination of drugs with the same characteristics (as considered in the previous section), the ratio that minimises the concentration required for almost certain treatment success is not either mono-therapy or 50:50, but can be something in between. Yet, in many cases, the optimal ratio both in terms of maximising *P*_success_ for a given concentration and in terms of minimising the concentration required for *P*_success_ *≥* 99% is still 50:50 or close to it. In cases where it is not, the mistake made by administering drugs at equal doses seems to be small (although it is unclear what should be considered ‘small’ here, given that the lives of patients may depend on it). In contrast, in cases where an unequal ratio is optimal (e.g. panel L), deviating from the optimal ratio in the other direction (i.e. giving even more of the high-dosed drug) can be detrimental, i.e. strongly reducing the probability of treatment success.

Overall, our results show that an equal distribution of the drug concentration amongst two drugs in combination might not always be the ideal drug ratio. While it is mostly the best strategy when the two drugs have the same drug characteristics and generally for low mutation probabilities, an unequal ratio is often beneficial when the two drugs vary in their Hill coefficients or modes of action, and the mutation probability is high. Nevertheless, the examples considered here suggest that, when information is limited, a 50:50 ratio is a good choice since it is usually not much worse than the best strategy (an exception are drugs with steep dose response curves and high mutation probabilities, as discussed before).

### 3.5 Comparison of combinations with more than two drugs

So far, we have focused on combinations with only two drugs, but what happens if we increase the number of drugs? In this section, we consider combinations of up to seven drugs to generalize our results from section 3.2. All drugs in combination have the same dose-response curve and a bactericidal mode of action, killing independent of replication (CK). The total drug dose is split equally among them. We display the results for different Hill coefficients. Since more drugs are expected to be especially important when resistance pre-exists, we show those results in the main text. Varying the modes of action does not change the results discussed in the following (figures S22 and S23).

#### Bliss Independence and Loewe additivity agree: more is better when dose-response curves are shallow

Considering drugs whose joint effect is described by Bliss independence leads to the same conclusion as for drugs with Loewe additivity if the drugs’ Hill coefficients *κ*_*i*_ *≤* 1: Increasing the number of drugs increases the probability of treatment success, even though the individual drug concentrations are reduced more with every drug added (figure 7A-C and G-I, figure S24). Yet, from a certain number of drugs, the benefit of adding more drugs becomes small. The same is found if we only account for the *de novo* evolution of resistance, except that a two-drug treatment is then still much better than mono-therapy even at high mutation probabilities (figure S25). The increase in treatment efficiency when increasing the number of drugs can be explained similarly to the results comparing the two-drug combination and mono-therapy (see section 3.2). For each type, the net growth rate decreases with the number of drugs (see figure S8 for the net growth rates of susceptible cells, single and double mutants). The more drugs, the more mutations are required for resistance and the less likely resistant types are generated.

**Figure 7.**
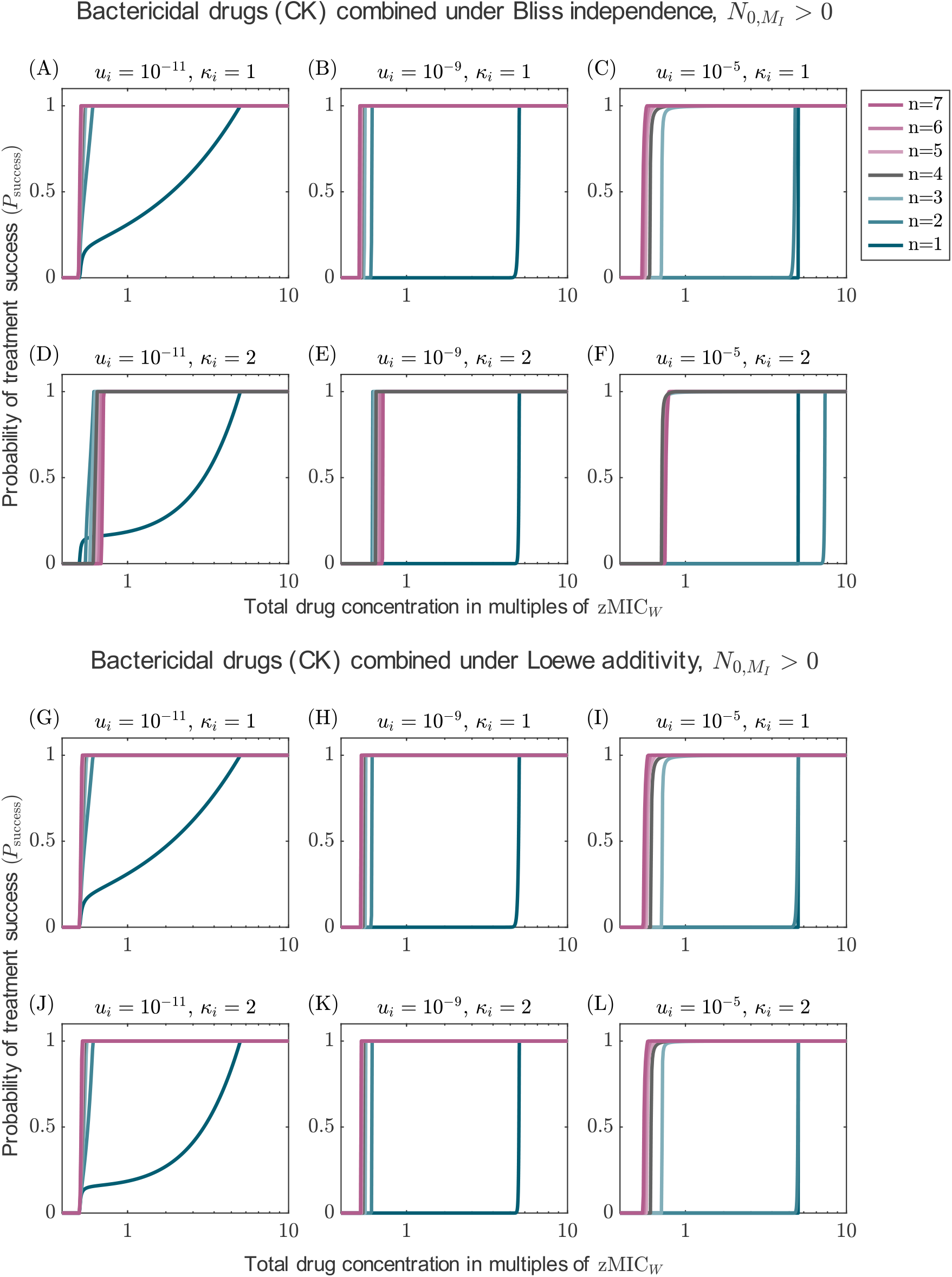
Probability of treatment success for combination with up to seven drugs for Bliss independence and Loewe additivity, accounting for pre-existing resistance mutations. The results are displayed for different mutation probabilities (columns) and different Hill coefficients (rows). All drugs are bactericidal, killing independent of cell replication (CK).

#### Bliss Independence and Loewe additivity disagree: more is not always better when dose-response curves are steep

While the results for drugs displaying Loewe additivity do not change for drugs with steep dose-response curves, figure 7D-F show that for Bliss independence, more drugs are not always better. This can be explained in the same way as for comparing two-drug treatments and mono-therapy in section 3.2: The wild-type gets better suppressed by treatments with fewer drugs. The same holds for the type resistant to the respective treatment (single mutants get better suppressed by mono-therapy than double mutants by two-drug treatments, etc.). However, any resistant type is controlled better by adding another drug (e.g., single mutants are better controlled by multi-drug treatment than by mono-therapy). Which effect dominates varies with the drug concentration, the mutation probability during treatment, and the pre-existence of mutants. Accordingly, the optimal number of drugs depends on the conditions. As already observed in section 3.2, two-drug treatment performs worse than mono-therapy if the mutation probability is high and pre-existence is considered (panel F). Here, adding a third drug provides a huge benefit since – while double mutants are frequent – triple mutants are not.

As a rule of thumb we can formulate that increasing the number of drugs increases the probability of treatment success up to some point, but then the benefit decreases or – in some cases – even turns into a disadvantage. Overall, our results suggest that, when mutants appear frequently and two-drug treatments are inefficient, combining three drugs might already provide a sufficiently large benefit compared to combinations with two drugs.

## 4 Discussion

How do we best design combination treatments to limit the evolution of drug resistance? To answer this question, we set up a stochastic pharmacodynamic model for treatment comparison and explored how drug characteristics and dosing strategies affect resistance evolution. To understand where the benefits of a certain treatment come from, we look at two factors influencing the evolution of resistance: the number of mutants generated *de novo* and the establishment probability of the mutants, i.e. the treatments’s ability to prevent mutants from arising and to control the growth of mutants once present.

### The effect of the mode of action of drugs on the evolution of resistance

For mono-therapy, we found that bacteriostatic drugs are better at limiting the *de novo* evolution of resistance than bactericidal drugs, provided the effect on the net growth rate is the same. Under this condition, bacteriostatic drugs are better at preventing mutants from arising than bactericidal drugs but result in a higher establishment probability. Bactericidal drugs only become advantageous when mutants pre-exist or for large enough bactericidal effects.

Only recently, stochastic models started to compare how the type of drug effect – reducing the replication rate or increasing the death rate – affects the evolution of resistance (Marrec & Bitbol, 2020; Czuppon et al., 2023; Raatz & Traulsen, 2023). In a pharmacodynamic model similar to ours, Czuppon et al. (2023) observe higher establishment probabilities of resistant mutants for treatments with bactericidal than with bacteriostatic drugs, which is opposite to what we find. The reason for this difference is that Czuppon et al. (2023), unlike us, include strain competition (assuming that non-replicating cells are still metabolically active); this means that growth of resistant strains is suppressed by competition with the sensitive strain for treatments with bacteriostatic drugs. Whether a drug affects the replication or the death rate is also of importance in more complicated ecological or genetic settings, e.g. it affects the rate of resistance evolution in fluctuating environments (Marrec & Bitbol, 2020) or the number of lineages that evolve during an adaptive walk through the trait space (Raatz & Traulsen, 2023).

Overall, the results highlight the importance of differentiating between drug effects on birth and death rates and accounting for stochastic population dynamics rather than assuming a deterministic net growth rate in which bacteriostatic or bactericidal drugs are implemented by simply varying the size of the drug effect (i.e. *ψ*_min_). Coates et al. (2018) demonstrate that the extinction probability predicted by a birth-death process (with experimentally measured rates) is consistent with the experimentally determined probability of stochastic extinction of susceptible cells (basically ‘proving’ our Eq. (A11) but for the wild type). This nicely shows that the mathematical approach of birth-death processes is suitable to describe stochastic bacterial dynamics. Extending the work by Coates et al. (2018) for more drugs and environmental conditions would help to improve model implementations and predictions on the efficiency of treatments using drugs with different modes of action.

### Comparing mono-therapy with a two-drug combination

In most cases, two-drug combinations, administered at equal drug ratios, are better than mono-therapy. Exceptions are found for drugs with steep dose-response curves (i.e. with a Hill coefficient larger than one) that combine according to Bliss independence. For those drugs, mono-therapy controls susceptible and fully resistant types better than the combination and is therefore sometimes the better treatment choice, especially when mutation rates are high and mutants pre-exist. A pharmacokinetic-pharmacodynamic model for sequential therapy, where drug doses overlap at each administration, previously found that the same effect turns the maximum cycling frequency sub-optimal for such drugs (Nyhoegen & Uecker, 2023).

*In vitro* studies generally confirm that combination treatment can reduce resistance evolution compared to mono-therapy (Hegreness et al., 2008; Munck et al., 2014; Barbosa et al., 2018; Jahn et al., 2021; Angst et al., 2021), but show that the ability to reduce resistance evolution is influenced by drug-drug interactions (Hegreness et al., 2008; Barbosa et al., 2018) or collateral effects (Munck et al., 2014; Barbosa et al., 2018). In clinical studies, resistance evolution is rarely considered as an endpoint. Among the few studies that do so, some observed a decrease in resistance evolution when using antibiotic combinations rather than mono-therapies; others, however, find no difference, leaving the comparison inconclusive for the clinic (Tamma et al., 2012). Similarly, a recent meta-analysis of 29 studies assessing the effect of combination therapy on the within-patient evolution of resistance in comparison to mono-therapy did not identify a general effect (Siedentop et al., 2023). However, the authors further showed that most of the clinical trails did not possess sufficient statistical power, highlighting the need for much larger clinical studies to fill this important knowledge gap. Mathematical models which incorporate important aspects of patient treatment such as the pharmacokinetics of the drugs might help to understand how our findings and those from *in vitro* studies translate to the clinic.

We further extended our comparison to combinations of drugs at unequal drug ratios. While equal drug ratios are most often beneficial for low mutation rates, unequal drug ratios can be optimal at high mutation rates, especially when combining drugs with different characteristics. However, a 50:50 ratio is usually not much worse. To our knowledge, the effect of different drug ratios on the evolution of resistance has not yet been systematically studied. We only considered a limited set of examples here, and an in-depth exploration of the parameter space is required to further assess the importance of the drug ratio. As it is impossible to achieve a specific drug ratio within the patient, such an analysis needs to consider the errors made by deviating from the optimal ratio.

### Combinations of drugs with different modes of action

While combination treatments, irrespective of the modes of action, are usually better at limiting resistance evolution than mono-therapy, not all combinations are equally good. For all cases considered here, the combination of a bacteriostatic drug with a bactericidal drug acting independent of replication was the best or one of the best choices. Combining instead a bacteriostatic drug with a bactericidal drug acting during replication led in most cases to the lowest success probability among the combination treatments included in the comparison (but the combination was still better than mono-therapy).

Whether bacteriostatic and bactericidal drugs should be combined, is also of interest simply with respect to their potency to clear susceptible cells. Coates et al. (2018) showed *in vitro* that adding a bacteriostatic drug to a bactericidal drug that acts on non-replicating cells can strongly increase stochastic extinction of susceptible cells. For bactericidal drugs that are only effective against replicating cells, in contrast, adding a bacteriostatic usually had no or even an adverse effect. Based on Loewe additivity, many combinations of bacteriostatic and bactericidal drugs are furthermore classified as antagonistic (Ocampo et al., 2014; Chandrasekaran et al., 2016) or even suppressive (Lázár et al., 2022). However, Ocampo et al. (2014) also found synergistic combinations of bacteriostatic and bactericidal drugs, especially for combinations including aminoglycosides, which have been found to kill non-replicating cells (McCall et al., 2019). Overall, combining bacteriostatic and bactericidal drugs can be a good strategy to limit the evolution of resistance. However, choosing such a combination requires a good understanding of how the drugs affect cellular processes and how these respective processes influence each other.

### Combinations with more than two drugs

Increasing the number of drugs often increases the treatment efficiency. However, the benefit of adding another drug decreases with an increasing number of drugs. For drugs displaying Bliss independence, an increase in the number of drugs can even decrease the treatment efficiency when the drugs have steep dose-response curves, as already discussed for two-drug treatments. In this case, our results suggest using a two-drug combination (or even mono-therapy) for very low and a combination of three or four drugs (depending on the mode of action) for large mutation probabilities. Increasing the number of drugs is a beneficial strategy for rapidly evolving strains. The current guidelines for the treatment of *M. tuberculosis* suggest using a four-drug combination in the first six months of treatment (WHO, 2022). Such multi-drug combinations are common for other rapidly evolving pathogens such as HIV and Malaria (Goldberg et al., 2012). The use of more than two drugs can even be necessary when only a small fraction of the cells in the population have an elevated mutation rate (i.e. mutator-strains): in a recent study combining laboratory experiments and modeling, Gifford et al. (2023) observed resistance evolution against the combination of rifampicin and nalidixic acid for a population of *E. coli* in which only 6% of the bacteria had an elevated mutation rate compared to the wild-type, while multi-drug resistance did not evolve in the absence of the mutator strain.

To provide a baseline comparison, we decided not to include drug-drug interactions in our analysis. Nevertheless, increasing the number of drugs can increase the frequency of drug-drug interactions (Katzir et al., 2019; Tekin et al., 2018). Hence, accounting for interactions – including possibly higher-order drug interactions – is a relevant next step.

### Contrasting the results obtained for Loewe additivity and Bliss independence

Whenever possible, we included results obtained for both additivity models, Loewe additivity and Bliss independence. Generally, the conclusions drawn from the two models are similar. However, for drugs with steep dose-response curves, the conclusions sometimes diverge due to the shift in MIC with Bliss independence, as discussed above. The need for higher drug doses with an increasing number of drugs aligns with findings from *in vitro* experiments (Russ & Kishony, 2018). This study concludes that Bliss independence can better predict the effect of multi-drug combinations than Loewe additivity (see also Katzir et al., 2019). Given the assumptions on the target sites made by the two models, this could be explained by the decreasing likelihood that all drugs target the same cellular site with an increasing number of drugs.

As mentioned in the method section, there are puzzles with implementing Bliss independence on rates. For bactericidal drugs, the common approach of summing up the death rates leads to an implausible high maximum effect, especially for large numbers of drugs. This approach as well as our implementations for the other modes of action further actually do not align with the original definition by Bliss (1939). How to translate Bliss independence to replication and death rates is not trivial and raises fundamental questions that we lack the space to address here. Modelling the drug effect explicitly through drug-target interactions (extending for example the work by Baeder et al. (2016)) might bring some clarity. Alternatively, modelers could work with specific experimentally measured dose response curves without caring how the combination is classified (e.g. Ankomah et al., 2013). For stochastic models, comparisons of time kill curves of drug combinations and mono-therapies (such as in Ankomah et al. (2013) and Yu et al. (2016)) would need to be extended by measurements of the replication and death rates of cells.

### Limitations and extensions

A caveat to our study is that – since we focused on a detailed understanding of the observed patterns – we only considered a limited set of parameters. While we varied several important parameters to test for robustness and many explanations seem intuitive, we cannot rule out that there may possibly be regions in the empirically relevant parameter range where results deviate. A study that focuses on a parameter sensitivity analysis would therefore be a highly valuable complement.

Apart from this general caveat, one of the major limitations of our work is that we only considered one resistance mutation for each drug, while in reality there are several or many mutations that can confer various levels of resistance. Generally, more mutations confer resistance to low than to high drug concentrations, which could impair the efficiency of multidrug low-dose treatments. Single resistance further evolved in our model with the same probability and resulted in the same increase in MIC for all drugs. The optimization with respect to the drug ratio might become more critical if resistance to one drug is more frequent or stronger than to the other. We did not include cross-resistance, which – depending on its strength – might make combination treatment lose its benefit over mono-therapy.

We only considered the acquisition of resistance through chromosomal mutations. Yet, resistance genes are often located on plasmids (Carattoli, 2013). Pathogenic bacteria could either carry a resistance plasmid to begin with or acquire it through horizontal gene transfer from commensal bacteria (Francino, 2016; San Millan, 2018) or other pathogens during a polymicrobial infection (Orazi & O’Toole, 2019). It is reasonable to assume that, similar to mutations, the acquisition rate is proportional to the population size; e.g. conjugative transfer is often modeled by a mass-action kinetics term (Levin et al., 1979). However, the rate depends on the dynamics of the donor population, which might be affected by the treatment too or interact with the recipient population. For cells whose growth is inhibited by a bacteriostatic drug, resistance on a newly acquired plasmid might not be expressed as for example assumed in the model by Willms et al. (2006). Importantly, the acquisition of a multidrug resistance plasmid would lead to resistance to multiple drugs in one step. There are numerous further important differences to chromosomal resistance, e.g. the risk of segregative loss of incompatible plasmids (Ishii et al., 1978; Novick & Hoppensteadt, 1978; Cullum & Broda, 1979) or the effect of conjugation on the establishment of the resistance plasmid (Novozhilov et al., 2005; Tazzyman & Bonhoeffer, 2013, 2014; Geoffroy & Uecker, 2023). Our model is thus most relevant for treatments with antibiotics where resistance relies on mutations and cannot be acquired through plasmids, but it also applies to other antibiotics as long as no resistance plasmid is available to enter the population. For reviews on modeling plasmid dynamics and an introduction to plasmid biology for modelers with modeling examples, we refer to Hernández-Beltrán et al. (2021) and Dewan and Uecker (2023).

Concerning the modes of action, it has been criticized that drugs cannot be strictly classified as either bacteriostatic or bactericidal (Pankey & Sabath, 2004; Wald-Dickler et al., 2018) and that furthermore, the static or cidal activity might depend on the antibiotic concentration (Pankey & Sabath, 2004). In a model that accounts for that, the drug would thus need to independently act on both rates, possibly with primarily bacteriostatic effects at low and primarily bactericidal effects at high concentrations. We also neglect many aspects of bacterial growth, especially competition, which has repeatedly been found to affect the evolution of single and multi-drug resistance (e.g. Pena-Miller et al., 2013; Berríos-Caro et al., 2021; Czuppon et al., 2023).

We restricted our analysis to constant drug concentrations. In the patient’s body, the drug concentration declines over time due to pharmacokinetic processes (Levison & Levison, 2009), which could reduce the efficiency of a combination compared to mono-therapy when administered at the same total drug dose. In addition, the pharmacokinetics of the drugs could vary, for example, in the elimination or penetration rate; hence, besides the concentration, the drug ratio could change with time. In extreme cases, this might lead to temporal or spatial mono-therapy, which can facilitate resistance evolution (Moreno-Gamez et al., 2015). Last, it would be important to also consider clinically relevant endpoints other than resistance evolution, such as the time until the infection is cleared. Here could lay a potential advantage of bactericidal over bacteriostatic drugs, as death rates can be much higher and hence eradication faster.

## Conclusion

With our study, we step by step developed a set of guiding principles of how to design combination treatments. We hope that it is the starting point for future studies that assess the robustness of the conclusions for multiple mutations with different resistance levels, unequal mutation probabilities for the two drugs, pharmacokinetic processes, and other factors, and extend the model by drug-drug interactions. Our study further highlights the need to better link the traditional phenomenological definitions of drug independence at the population level and the microscopic dynamics at the level of cells. A model that is consistent between the two levels would generate a common framework for mathematical modeling, experiments on single-cell dynamics, and traditional experimental measurements at the population level and would thus allow for a much better integration of modeling work and empirical results.

## Supporting information

Additional information for the methods and results section of the main paper.

## Data availability

All code used to generate the data displayed in the figures is available at https://gitlab.gwdg.de/nyhoegen/the-many-dimensions-of-combination-therapy-supplementary-code.

## Acknowledgments

We thank Roland Regös for sharing his insights on drug additivity models and pharmaco-dynamics. We further thank all members of the Research group Stochastic Evolutionary Dynamics (MPI) and the Research group Theoretical Biology (ETH) for their support and helpful discussions.

## Appendix

### A Numbers of cells of each type in mutation-selection balance

We derive the frequencies of susceptible and resistant bacteria in mutation-selection balance in the absence of drug based on the deterministic version of our model. Given for example treatment with two drugs, resistance can pre-exist against either or both of the drugs, hence, the population can consist of wild-type cells (*W*), single mutants (*M*_1_ and *M*_2_) and double mutants (*M*_1,2_). The differential equation system describing the dynamics of this population is given by:

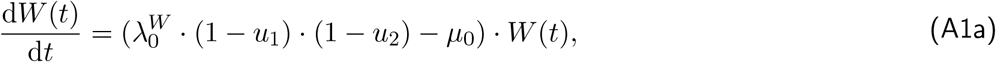

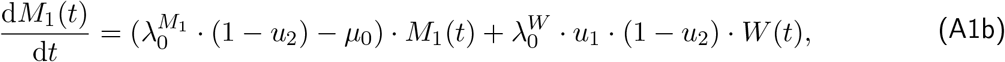

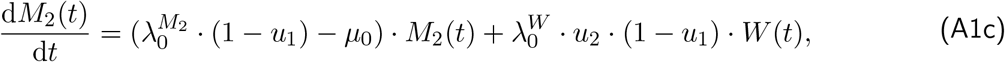

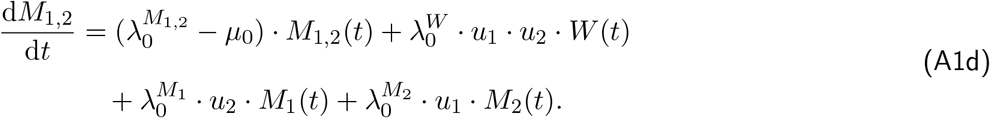

To obtain the frequencies of these cells at mutation selection balance, we solve the differential equation system over a long time period starting with one cell in MATLAB (ode45) until the relative frequencies have converged to their values in mutation-selection balance. The system (A1) describes an exponentially growing population. We could have introduced a carrying capacity to limit population growth, but the simple exponential model is sufficient to determine the mutation-selection balance.

For the parameter values of *λ*_0_ = 0.7, *μ*_0_ = 0.1 and *γ* = 0.1 we derive the following numbers of mutants pre-existing in a population of *N*_0_ = 10^10^ cells, considering up to five drugs (see table 4). The relative frequency of mutants that are resistant to more than two drugs is never above 1*/*10^10^, i.e. less than one cell; we consider these mutants absent, and the corresponding values are not displayed.

**Table 4:** Numbers of mutants pre-existing in a population of *N*_0_ = 10^10^ cells at mutation-selection balance for resistance against up to five drugs and for different mutation probabilities.

| Number of drugs | $u_i = 10^{-11}$ | $u_i = 10^{-9}$ | $u_i = 10^{-5}$ |
| --- | --- | --- | --- |
| 1 | $N_{0,M_1} = 1$ | $N_{0,M_1} = 100$ | $N_{0,M_1} = 1000000$ |
| 2 | $N_{0,M_1} = N_{0,M_2} = 1$<br>$N_{0,M_{1,2}} = 0$ | $N_{0,M_1} = N_{0,M_2} = 100$<br>$N_{0,M_{1,2}} = 0$ | $N_{0,M_1} = N_{0,M_2} = 999900$<br>$N_{0,M_{1,2}} = 100$ |
| 3,<br>$i, j \in \{1, 2, 3\}$ | $N_{0,M_i} = 1$<br>$N_{0,M_{i,j}} = 0$ | $N_{0,M_i} = 100$<br>$N_{0,M_{i,j}} = 0$ | $N_{0,M_i} = 999800$<br>$N_{0,M_{i,j}} = 141$ |
| 4,<br>$i, j \in \{1, 2, 3, 4\}$ | $N_{0,M_i} = 1$<br>$N_{0,M_{i,j}} = 0$ | $N_{0,M_i} = 100$<br>$N_{0,M_{i,j}} = 0$ | $N_{0,M_i} = 999700$<br>$N_{0,M_{i,j}} = 141$ |
| 5,<br>$i, j \in \{1, 2, 3, 4, 5\}$ | $N_{0,M_i} = 1$<br>$N_{0,M_{i,j}} = 0$ | $N_{0,M_i} = 100$<br>$N_{0,M_{i,j}} = 0$ | $N_{0,M_i} = 999610$<br>$N_{0,M_{i,j}} = 141$ |

### B Comparison of our model to Bliss’ original definition

As discussed in the main text, Bliss independence was originally defined through the proportions of a population killed by the respective treatments (Bliss, 1939). Baeder et al. (2016) derived mathematically how the joint drug effect of two drugs on the net growth rate can be calculated under Bliss independence. The joint effect is then given by the addition of the individual drug effects, which we use here for the combined drug effect if drugs affect the death rate. For our analysis we derive the extinction probability of a single wild-type cell (see section C), which is the counter-event to survival. Ignoring mutation, this probability is given by 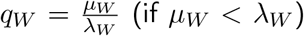, where *μ*_*W*_ and *λ*_*W*_ depend on the treatment. Based on this probability, we can compare our implementation with Bliss’ original definition. For a large number of cells, the probability *q*_*X*_ corresponds to the proportion of cells that would be killed (assuming we wait long enough). The (expected) proportion surviving treatment with drug *i* is given by one minus the extinction probability:

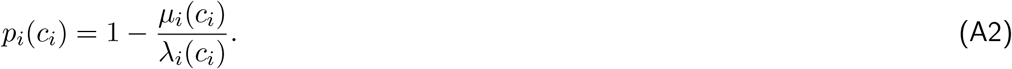

For the combination of two bactericidal drugs (CK), Bliss’ definition of the joint drug effect under stochastic independence (equation (8)) results in

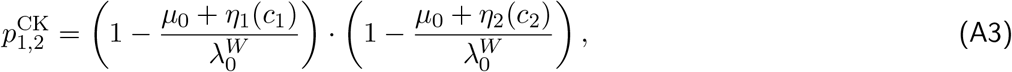

while the definition of Bliss independence on rates results in

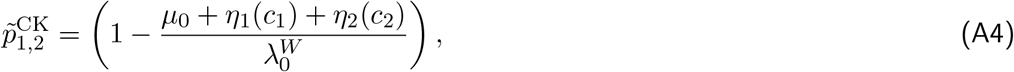

which thus does not fit the original definition. Note that we look here at the fates of cells (survival or extinction of the lineage) at infinite times; for a comparison at finite times, we would need to determine the probability that a cell lineage is extinct by time *t*.

The results are also different for the other two modes of action, although the joint inhibition of two bacteriostatic drugs is defined based on stochastic independence.

Calculating the probability of surviving the treatment with two bacteriostatic drugs results according to the original definition of Bliss independence in

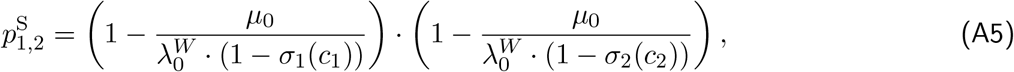

while we get

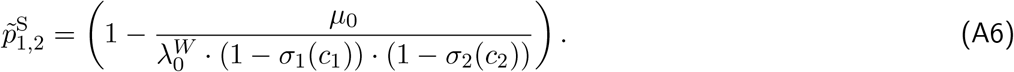

Similarly, Bliss’ definition of surviving the treatment with two bactericidal drugs (CR) results in

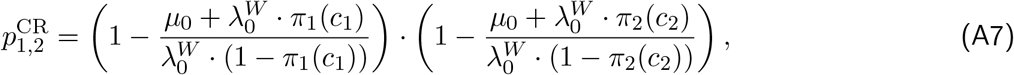

while we get

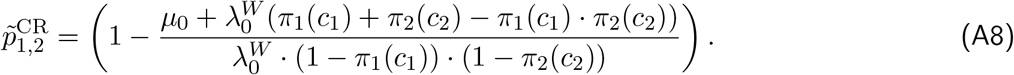

### C Exact calculation of the extinction probability

In the main text, we described the population dynamics with a time-continuous process. To determine the extinction probabilities, we transform the time-continuous birth-death process into a multi-type Galton-Watson process in discrete time. For this, we jump to the next event in the ‘life’ of each cell, which is either replication or death. If a lineage goes extinct, it will do so with the same probability as in the time-continuous process.

The type-specific replication and death probabilities, *ρ*_*X*_(*c*) and *δ*_*X*_(*c*), are calculated from the type-specific rates *λ*_*X*_(*c*) and *μ*_*X*_(*c*), which depend on the treatment, as discussed in the sections 2.2 and 2.3, and are given by

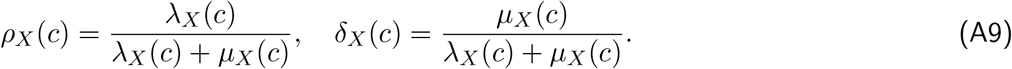

In the following we will first present an intuitive derivation of the extinction probabilities for the example of mono-therapy, before we provide the formal derivation for the treatment with an arbitrary number of drugs. Let us first consider the fate of a lineage starting from a mutant cell. The lineage goes extinct, if either the initial cell dies, which occurs with probability *δ*_*M*_ (*c*), or if the cell replicates (with probability *ρ*_*M*_ (*c*)) and both lineages founded by the two daughter cells go extinct. Due to the independence of lineages, the probability that both daughter cells fail to establish a long-term lineage is the product of the single probabilities. It hence holds that

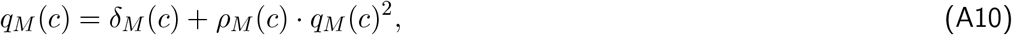

which yields

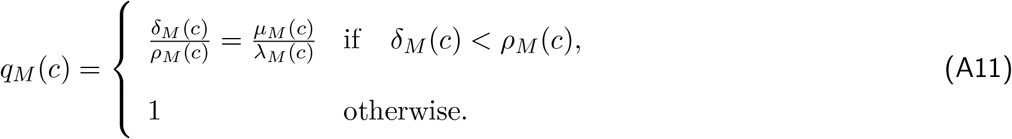

(Which root to choose follows from general branching process theory (Sewastjanow, 1974, p. 31). But it is also intuitive: if the probability of cell death is higher than or equal to that of cell birth, the lineage cannot persist indefinitely. If it is smaller, there is a positive chance that the lineage grows.)

The extinction probability of a lineage starting with a wild-type cell *q*_*W*_ can be derived in the same form, except that during replication, the cell might mutate. The lineage starting with a mutated daughter cell goes extinct with probability *q*_*M*_ as derived above, and we have

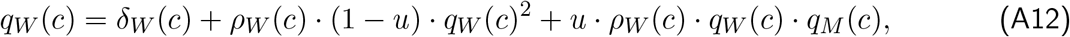

which can be solved to obtain *q*_*W*_ (*c*). The procedure can be generalized in a straightforward way to treatments with more drugs. Generally, since we assume that cells can gain but not loose mutations, the extinction probability of a type can always be determined recursively from those with more mutations.

We now proceed to give the formal derivation of the extinction probabilities *q*_*X*_ (*c*) founded by a single cell of type *X*. For notational simplicity we drop the dependency on the concentration *c*. To calculate *q*_*X*_, we consider the probability generating function corresponding to the ‘offspring’ distribution in the multi-type Galton-Watson process. For a treatment with *n* drugs, there are 2^*n*^ possible cell types in the population, and the probability generating function **f** (**s**) is thus a vector of length 2^*n*^, and the argument **s** is a vector of the same length.

The component of **f** (**s**) that corresponds to cell type *X* is given by

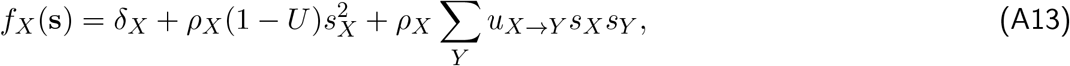

where *s*_*X*_ and *s*_*Y*_ are the components of **s** corresponding to types *X* and *Y, u*_*X→Y*_ is the probability that a daughter cell of type *X* is of type *Y*, and 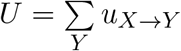. The probability *u*_*X→Y*_ is zero unless type *Y* carries additional mutations with respect to type *X*.

As an example, we explicitly show the probability generating function for *n* = 2. In a treatment with two drugs, there are four possible types – the fully susceptible wild type (*W*), the fully resistant double mutant (*M*_1,2_), and the two single resistant types (*M*_1_ and *M*_2_). The probability generating function **f** (**s**) with 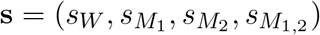 is given by the vector 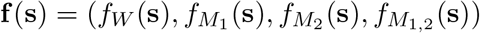with

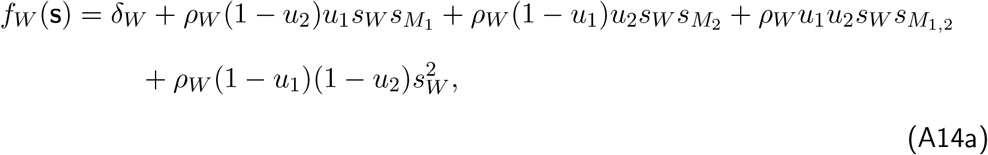

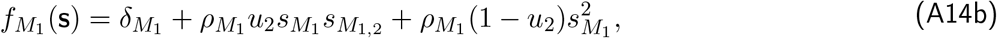

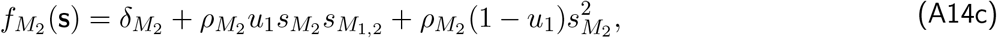

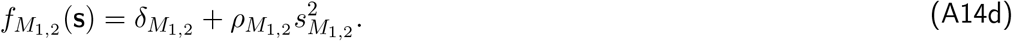

The vector of extinction probabilities **q** (e.g.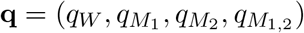) for treatment with two drugs) is the smallest fix point of the probability generation function (Sewastjanow, 1974, p. 115). One can convince oneself that the fix point equation **q** = **f** (**q**) corresponds to the equations that one obtains with the intuitive arguments in the main text.

### D Approximation of the probability of treatment success for a two-drug combination

In this section, we derive the approximation of *P*_success_ for a two-drug combination (drug 1 and drug 2), corresponding to approximation (14) for mono-therapy. Each term – number of new mutants and their establishment probabilities – needs to be calculated for each type that is resistant at a given concentration. We model the dynamics of all types that have a negative growth rate at that concentration deterministically and use this to obtain the number of new mutants resistant to the treatment. We then calculate their establishment probabilities without considering further mutations. I.e. in concentration ranges where a single mutant has a positive net growth rate, we ignore the possibility that the single mutant might go extinct but generate a double resistant type before doing so. I.e., we only consider two-step rescue via single mutants when single mutants cannot grow themselves. Hence, we split the concentration range into two parts. The first range includes all concentrations at which the single resistant types have a positive growth rate. For simplicity, we assume here that this range is the same for both single resistant types, which does not need to be true generally (in those cases the concentration range needs to be split into more parts). We formulate the establishment probabilities based on rates which follows from the transformation in equation (A9).

For the first drug range, the mean numbers of mutants generated from the susceptible type (mean of a Poisson distribution) and their establishment probabilities are – nearly identical to the approximations for mono-therapy – given by:

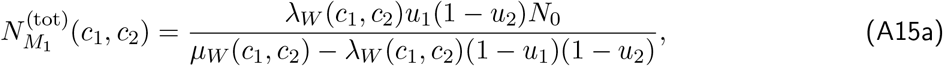

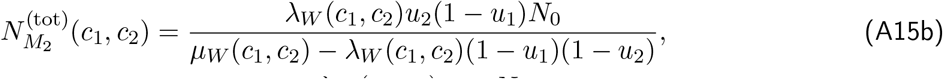

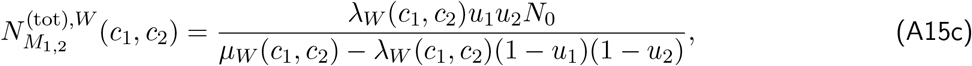

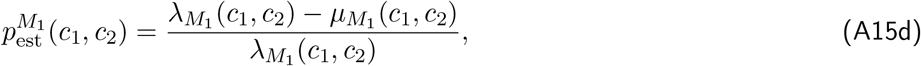

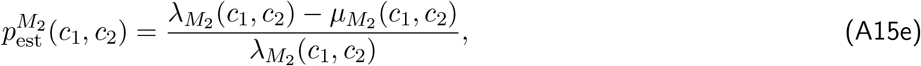

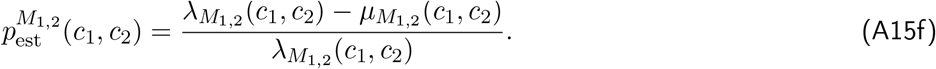

The probability of treatment success is then approximated by

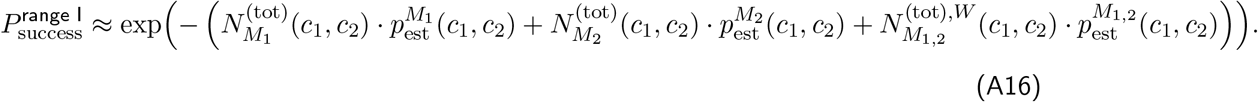

In the second drug range, we need to additionally consider the number of double mutants generated by the single resistant types (which are themselves doomed to extinction in this range). The number of single mutants generated by the wild type is again Poisson distributed by the mean given by the same expression as before (equation (A15)). We further describe the decay of theses single mutants deterministically as well (although they are rare), and the number of double mutants generated from each of these single mutants is again Poisson distributed with means

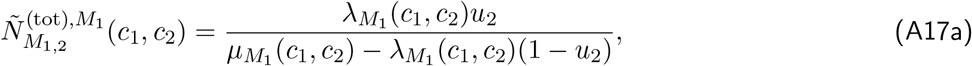

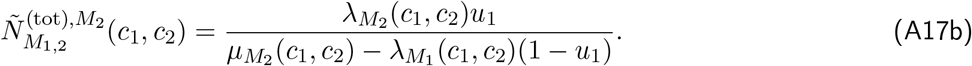

What is the probability distribution of the total number of double mutants generated in this way? Let us denote by *Y* the random variable describing the number of mutants resistant to drug 1 generated from the wild-type. *Y* is Poisson distributed with mean 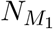, and the p.g.f. is thus given by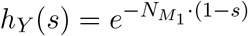. Let us denote by *X*_*j*_ the number of *successful* double mutants generated from the *j*th single mutant. *X*_*j*_ is Poisson distributed with mean 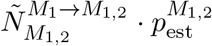, and the p.g.f. of is thus given by 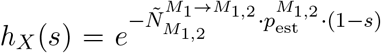. We are now interested in the p.g.f. of the random variable 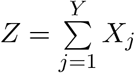, which is obtained by

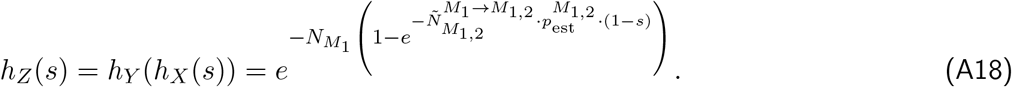

We further obtain for the probability that *Z* takes the value 0:

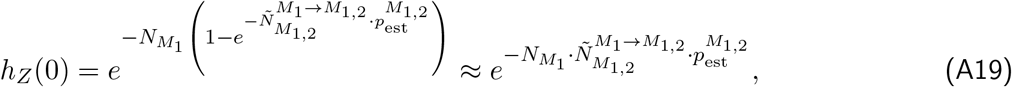

where the approximation assumes that 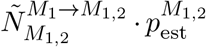 is small (which is a very reasonable assumption).

We now still need to account for (1) the same pathway to resistance via single mutants resistant to drug 2 and (2) double mutants that get directly generated from wild-type cells by two mutations. Setting

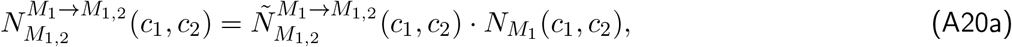

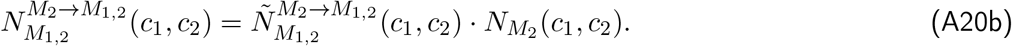

we obtain for the probability of treatment success

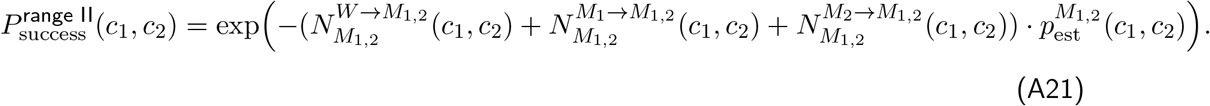

Given low mutation probabilities, terms (1 *− u*_1_) or (1 *− u*_2_) can be omitted since they are approximately one.

### E Targeting the replication rate, the death rate, or both: A comparison of mono-therapies

In section 3.1 in the main text, we investigated the effect of the mode of action of drugs in mono-therapy on the probability of treatment success. We observed that bacteriostatic drugs (S) are better than bactericidal drugs (CK and CR) and bactericidal drugs acting during replication (CR) are better than bactericidal drugs killing independent of replication (CK), given the dose-response curves are the same. In the following we will prove this mathematically.

Consider a drug where a fraction *α* of the drug effect *E*(*c*) is allocated to decreasing the replication rate and a fraction (1 *− α*) of the drug effect is allocated to increasing the death rate, i.e. *λ* = *λ*_0_ *− αE*(*c*) and *μ* = *μ*_0_ + (1 *− α*)*E*(*c*). For the S drug, we have *α* = 1; for the CK drug, we have *α* = 0; and for the CR drug, we have 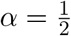 (see section 2.2).

For such a drug, the total number of new mutants that get generated during population decline is given by

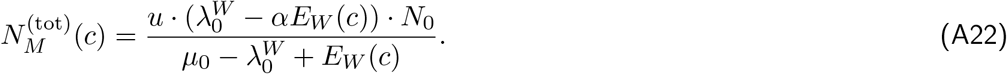

(We are omitting the term (1 *− u*) here since it is small.) This is a decreasing function in *α*, i.e. the more the drug acts on the replication rate, the fewer mutations are generated.

For the establishment probability of mutants, we have

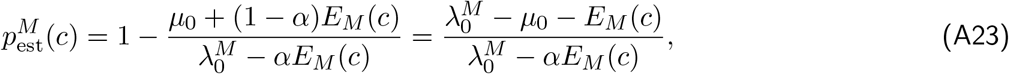

which is an increasing function in *α*. I.e., the more the drug acts on increasing the death rate, the lower the establishment probability of mutants.

What about the product of the two?

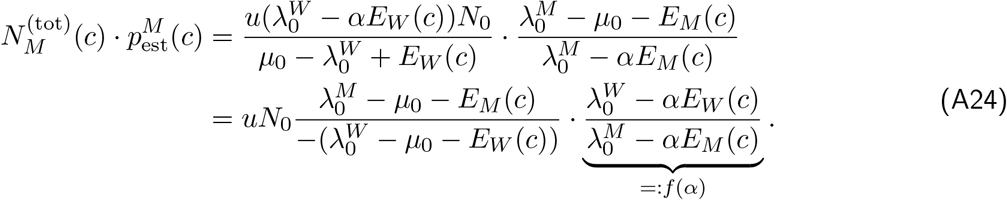

For the derivative of the function *f* (*α*), we obtain

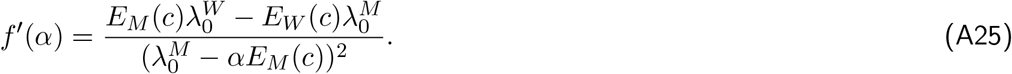

It holds:

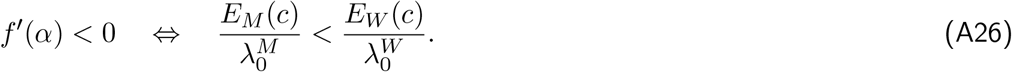

In the regime that we are interested in – the wild-type has a negative and the mutant a positive net growth rate –, it holds

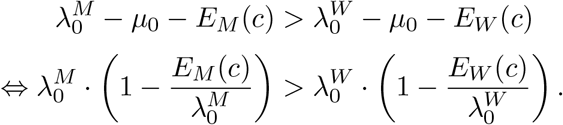

Since we assume that *λ*^*M*^ *≤ λ*^*W*^, it must especially hold that

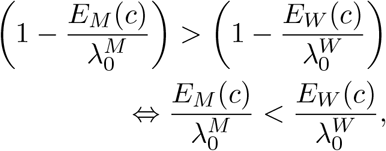

which shows that *f* ^*′*^(*α*) *<* 0. This in turn means that the product 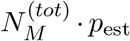 decreases with *α*. The smaller the product, the larger the probability of treatment success. Thus, the more drug effect is allocated to reducing the birth rate, the higher the probability of treatment success. This especially proves the order S *>* CR *>* CK.

