## Additional information for the methods and results section of the main paper. for "The many dimensions of combination therapy: How to combine antibiotics to limit resistance evolution"

### S1 Dose response curves for different drug combinations and cell types

In this section we provide additional figures that display the net growth rate function for different drug combinations and types.

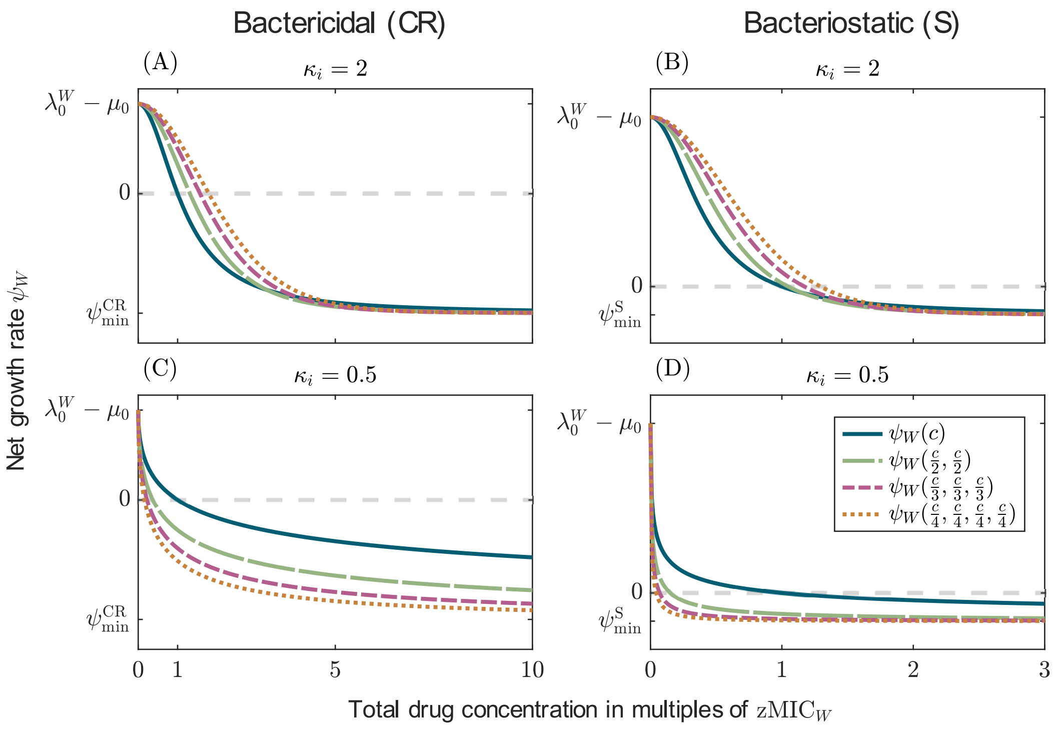


Figure S1: Net growth rate for treatments with up to four drugs under Bliss independence for bactericidal (CR) and bacteriostatic (S) drugs. The figure shows the net growth rate as a function of the total drug concentration for drugs with different Hill coefficients as indicated by the panel headers. The minimal growth rates are set to $\psi_{\min}^{\mathrm{CR}} = -0.8h^{-1}$ and $\psi_{\min}^{\mathrm{S}} = -0.1h^{-1}$.

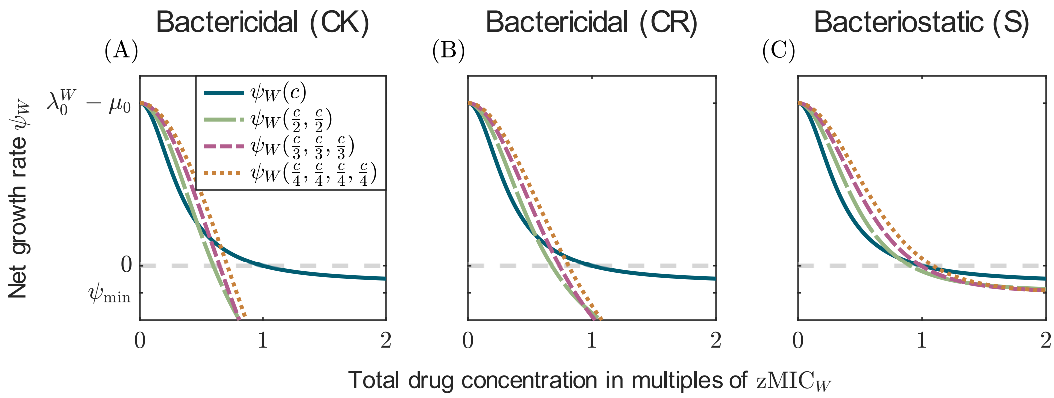


Figure S2: Net growth rate for treatments with up to four drugs under Bliss independence for bactericidal (CK and CR) and bacteriostatic (S) drugs. The figure shows the net growth rate as a function of the total drug concentration for drugs with Hill coefficients $\kappa_i = 2$. The minimal growth rates are set to $\psi_{\min} = -0.065h^{-1}$ for all drugs.

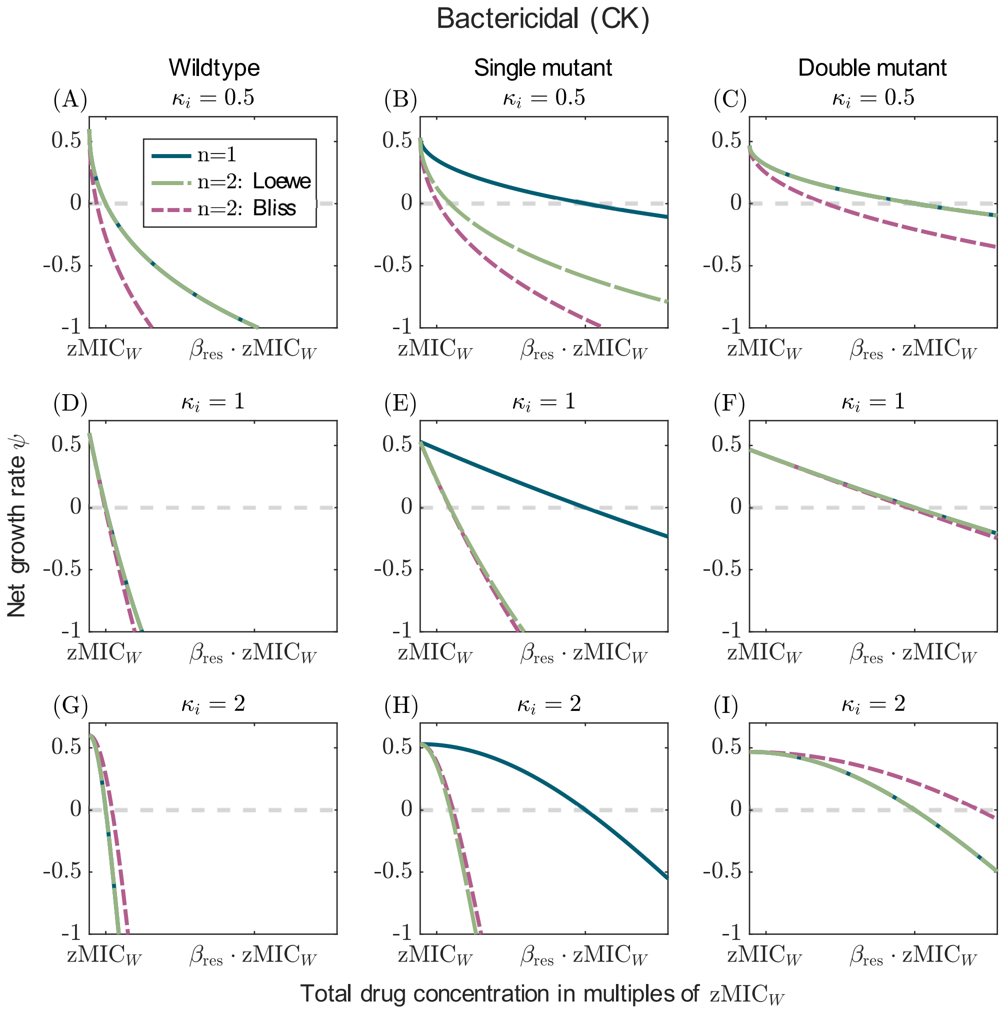


Figure S3: Comparison of the net growth rates of all types under mono-therapy and a two-drug combination of bactericidal drugs killing independent of replication (CK). Drugs can either display Bliss independence (pink line) or Loewe additivity (green line). The columns show the net growth rates of the wildtype, the single mutant (resistant to the first drug), and the double mutant (resistant to both drugs), respectively. The rows differ in the Hill coefficients of the drugs ($\kappa_i = 0.5$, $\kappa_i = 1$, and $\kappa_i = 2$ from top to bottom).

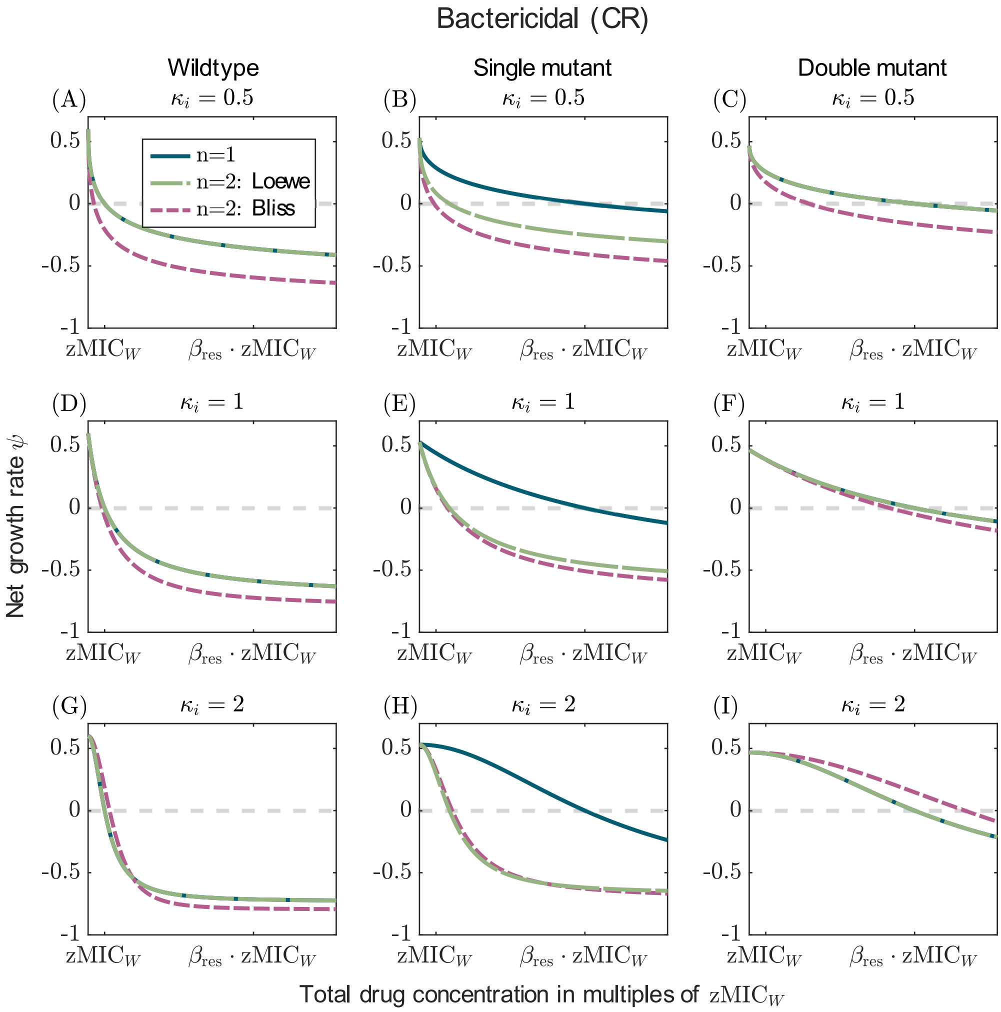


Figure S4: Comparison of the net growth rates of all types under mono-therapy and a two-drug combination of bactericidal drugs acting during replication (CR). Drugs can either display Bliss independence (pink line) or Loewe additivity (green line). The columns show the net growth rates of the wildtype, the single mutant (resistant to the first drug), and the double mutant (resistant to both drugs), respectively. The rows differ in the Hill coefficients of the drugs ($\kappa_i = 0.5$, $\kappa_i = 1$, and $\kappa_i = 2$ from top to bottom).

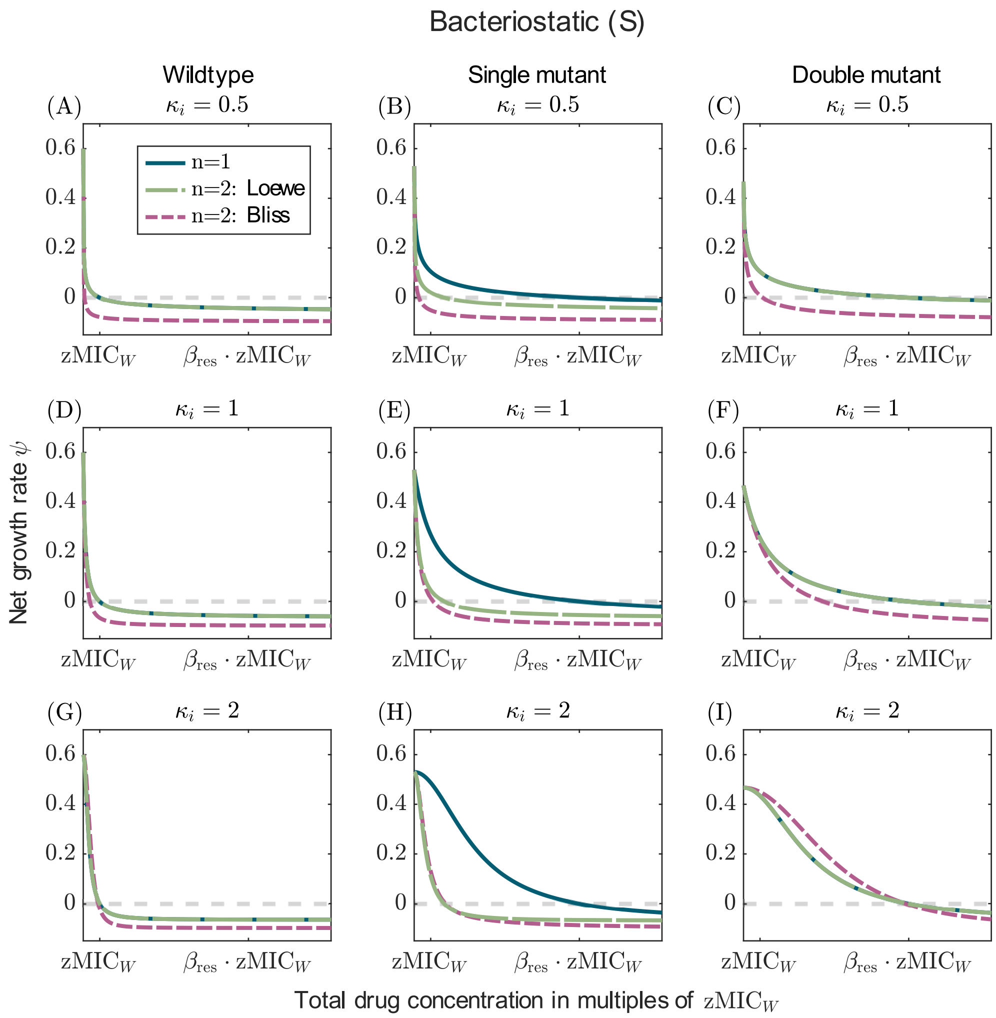


Figure S5: Comparison of the net growth rates of all types under mono-therapy and a two-drug combination of bacteriostatic drugs (S). Drugs can either display Bliss independence (pink line) or Loewe additivity (green line). The columns show the net growth rates of the wildtype, the single mutant (resistant to the first drug), and the double mutant (resistant to both drugs), respectively. The rows differ in the Hill coefficients of the drugs ($\kappa_i = 0.5$, $\kappa_i = 1$, and $\kappa_i = 2$ from top to bottom).

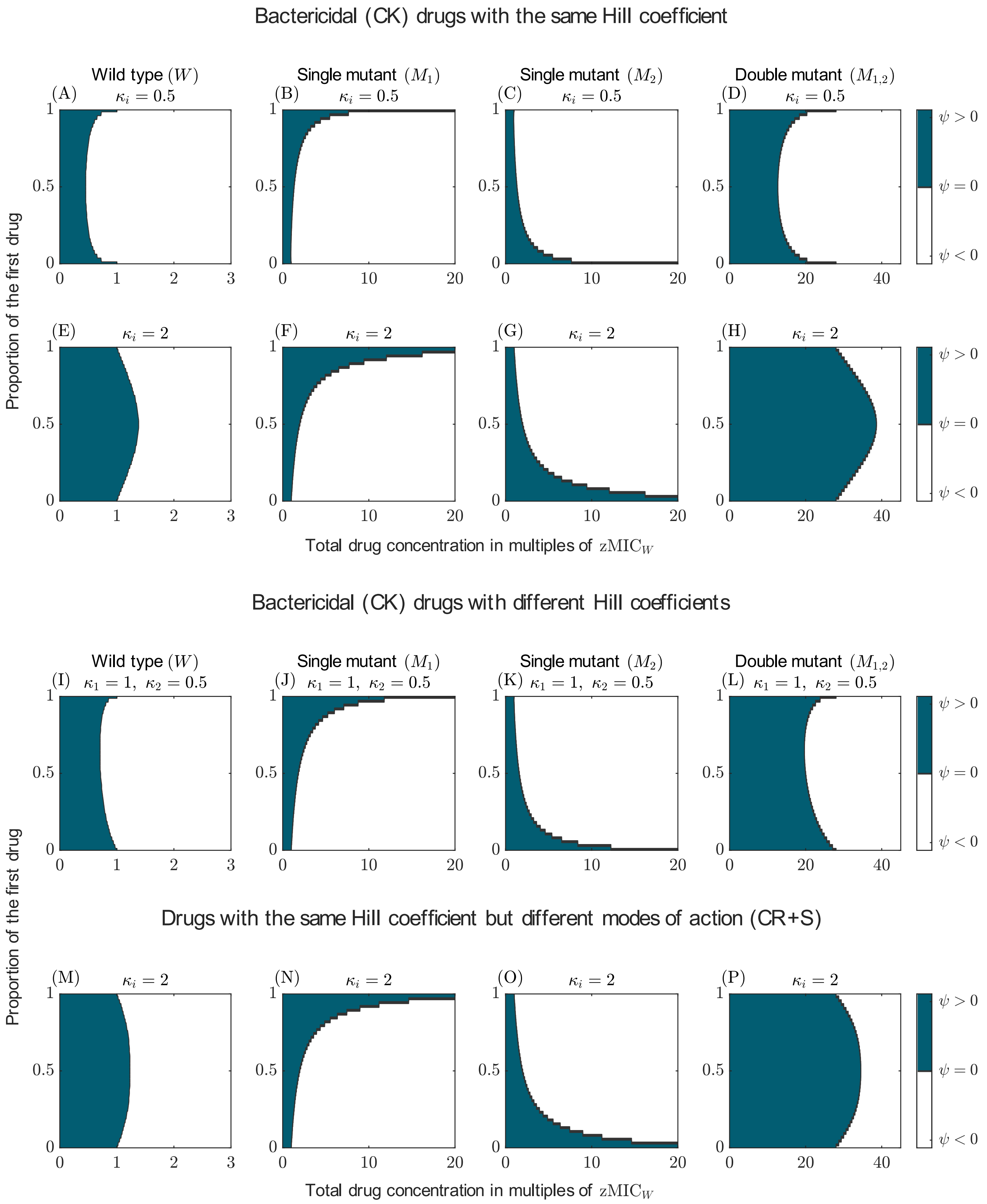


Figure S6: Ranges of positive and negative net growth for susceptible bacteria, single mutants, and double mutants for drug combination with two drugs administered at different drug ratios. The first column refers to the wild type, the second and third to the single mutants, and the fourth to the double mutant. The blue color indicates a positive and the withe color a negative net growth rate. The parameter values of the displayed treatments are the same as in main text figure 6.

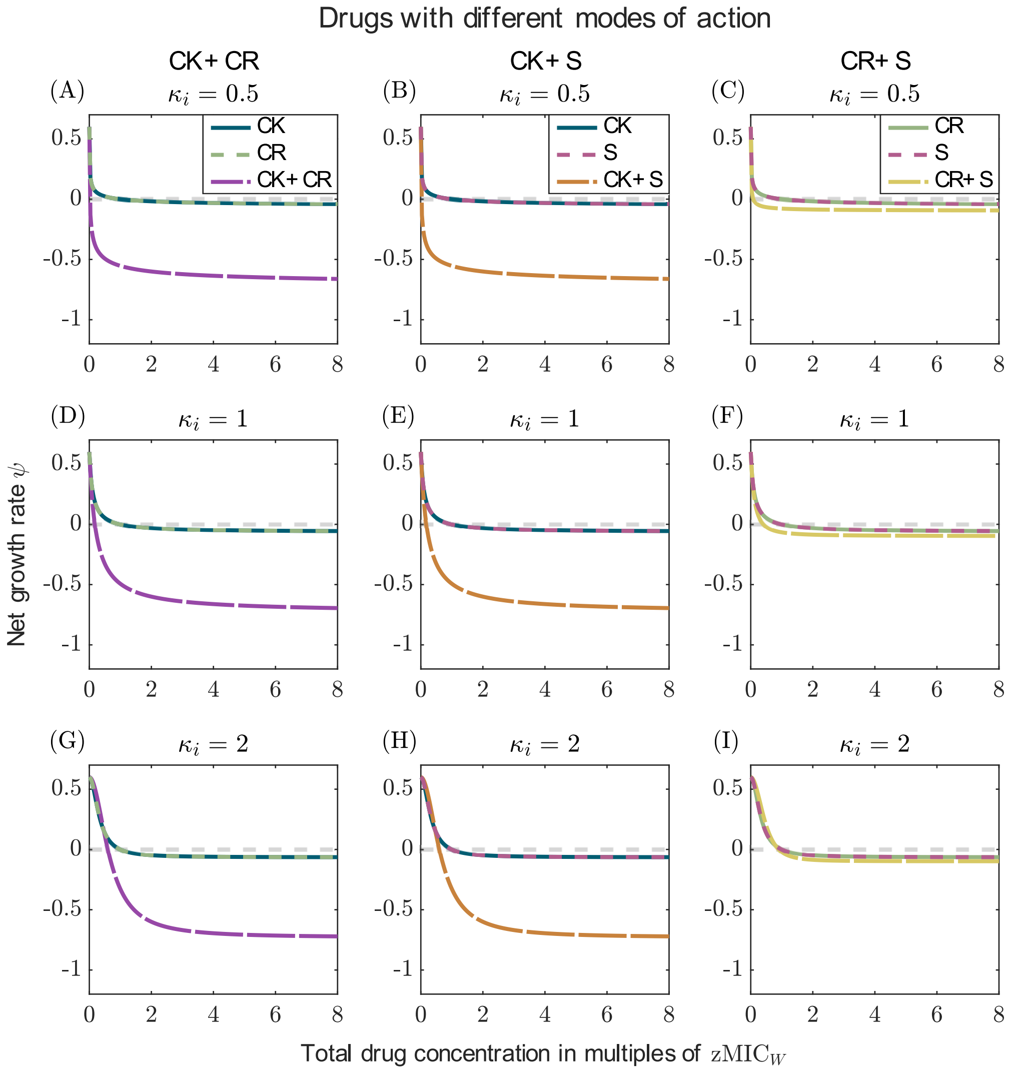


.

Figure S7: Net growth rates for treatments with drugs differing in their modes of action compared to the net growth rates under mono-therapy. The comparison is displayed for each combination of different modes of action (see column header) and for different Hill coefficients (rows). All drugs in mono-therapy have the same dose-response curves with $\psi_{\min} = -0.065^h - 1$.

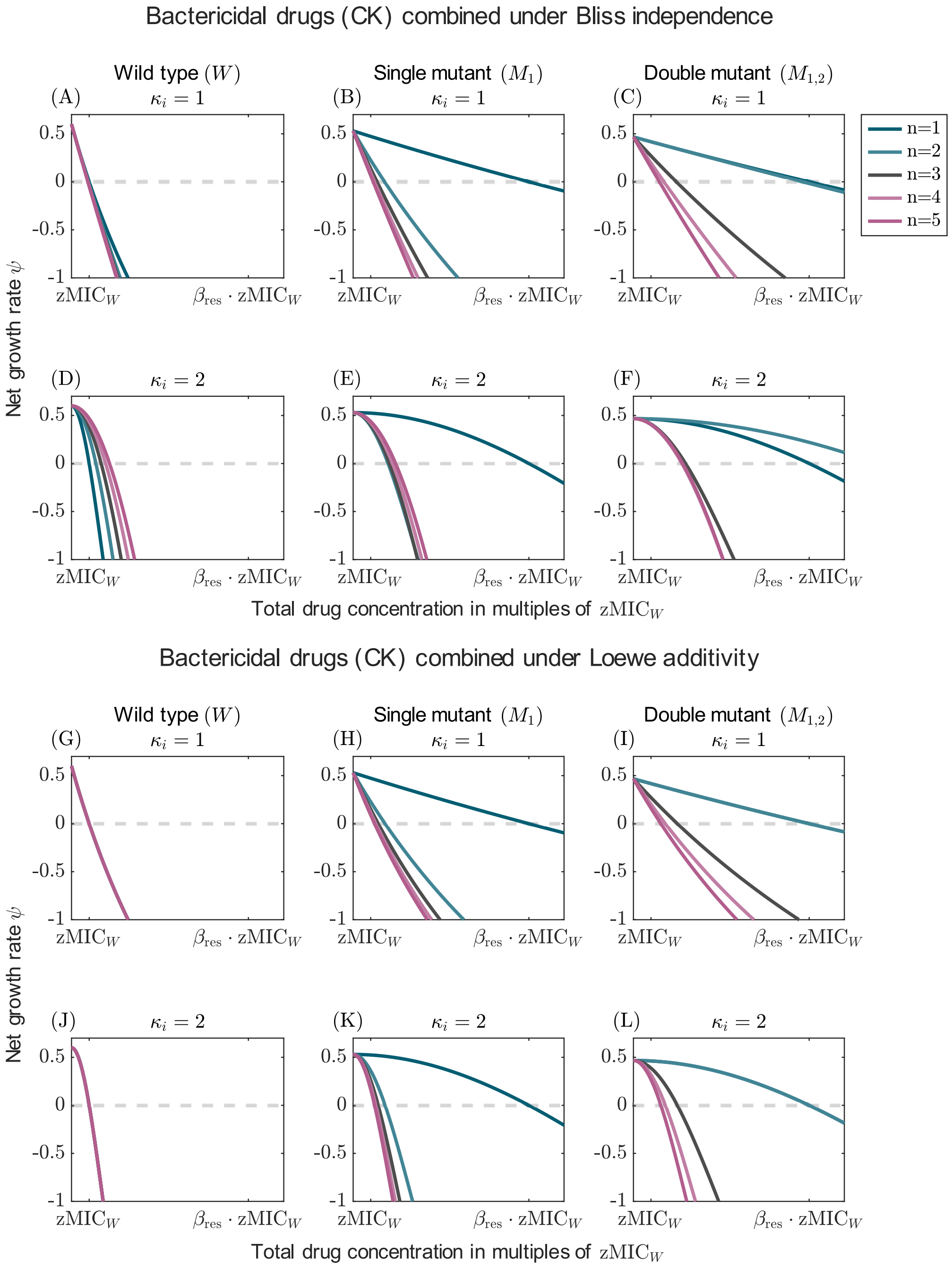


Figure S8: Net growth rates of susceptible bacteria, single mutants, and double mutants for treatments with up to five drugs. The net growth rate is displayed for the wild type (first column), the single mutant resistant to the first drug (second column), and the double mutant resistant to the first two drugs (third column). The upper part of the figure (panels A-F) shows results for drugs combining according to Bliss independence and the lower part (panels G-L) shows results for drugs combining according to Loewe additivity. The first row in each part is for drugs with $\kappa_i = 1$, the second row for drugs with $\kappa_i = 2$.

#### S2 Drug examples for different drug characteristics

In the analysis of our model, we accounted for different drug characteristics, such as different modes of action or different Hill coefficients of the drugs. The following table gives examples of drugs that display these characteristics. It further lists combinations of drugs with their respective modes of action that are recommended for the use in the clinic by official guidelines.

| **Mode of action** | Examples | Reference |
|---|---|---|
| Bactericidal[1] (CK) | Aminoglycosides | McCall et al. (2019) and Mutschler et al. (2005) |
| Bactericidal (CR) | $\beta$-lactam antibiotics<br>Ciprofloxacin and Rifampicin[1] | Tuomanen et al. (1986) and McCall et al. (2019)<br>Mutschler et al. (2005) |
| Bacteriostatic (S) | Macrolide, Tetracycline<br>Chloramphenicol, Ethambutol,<br>Folic acid antagonists,<br>Oxazolidinone, Nitrofurantoin | Mutschler et al. (2005) |
| CK + CR | Cefotaxime or Ceftazidime (CR)<br>+ Amikacin or Gentamycin (CK)<br>for sever urinary tract infections | WHO (2022) p. 477ff |
| S + CR | Ceftazidime (CR) + Azithromycin (S)<br>for severe entric fever | WHO (2022) p.192 |
| S + S | Trimethoprim + Sulfamethoxazole | Mutschler et al. (2005) |
| CR + CR | Vancomycin + Ceftazidime<br>for bacterial eye infections in adults | WHO (2022) p. 110 |
| **Hill coefficient** | Example | Reference |
| $\kappa < 1$ | Streptomycin, Clarithromycin<br>and Moxifloxacin (for *M. Marinum*) | Ankomah and Levin (2012) |
| $1 \leq \kappa < 2$ | Ciprofloxacin (for *E. coli, S. aureus*<br>and *N. gonorrhoea*)<br>Amikacin (for *M. marinum*)<br>Gentamicin (for *S. aureus*<br>and *N. gonorrhoea*)<br>Ceftriaxone (for *N. gonorrhoea*) | Regoes et al. (2004) and Ankomah et al. (2013)<br>Foerster et al. (2016)<br>Ankomah and Levin (2012)<br>Ankomah et al. (2013)<br>Foerster et al. (2016)<br>Foerster et al. (2016) |
| $\kappa > 2$ | Tobramycin (for *E. coli*)<br>Rifampicin (for *E. coli*<br>Azithromycin (for *N. gonorrhoea)* | Ankomah et al. (2013)<br>Regoes et al. (2004)<br>Foerster et al. (2016) |

Table S1: Examples of drugs for different drug characteristics. The table displays examples of drugs for different modes of action and ranges of Hill coefficients. It further lists examples for combinations of drugs with different modes of actions that are recommended for treatment by official guidelines. Note that Hill coefficients may vary across studies, even for the same bacterial strain.
[1]Rifampicin is able to kill non-replicating cells if the stasis is caused by the exposure to a bacteriostatic drug (McCall et al., 2019).

### S3 Comparison of mono-therapy and two-drug treatments for combinations of drugs with the same mode of action combined at a 50:50 ratio

In this section we display additional figures for the comparison of mono-therapy and two-drug combinations, assuming either Bliss independence or Loewe additivity. The results displayed in these figures extend the results displayed in main text figure 3 (see section 3.2).

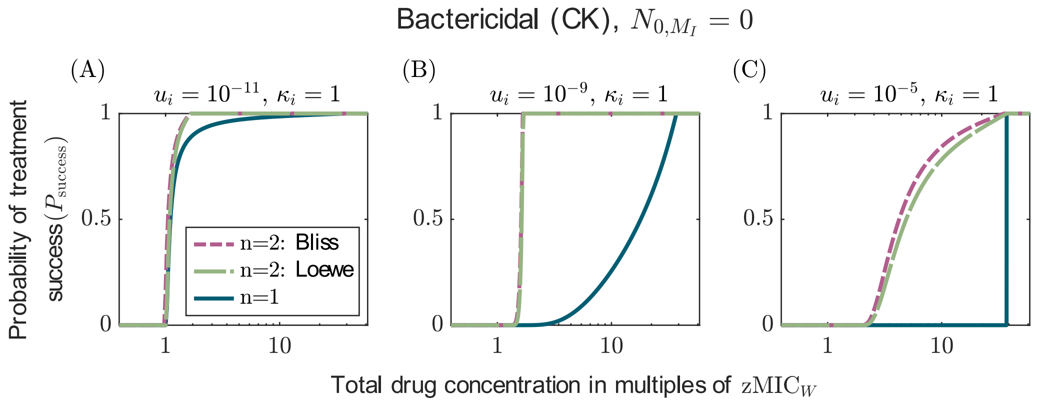


Figure S9: Comparison of the probabilities of treatment success under mono-therapy ($n = 1$) and a two-drug combination ($n = 2$) of drugs killing independent of replication (CK) for $\kappa_i = 1$. The figure corresponds to figure 3 with the difference that that it considers less steep pharmacodynamic curves.

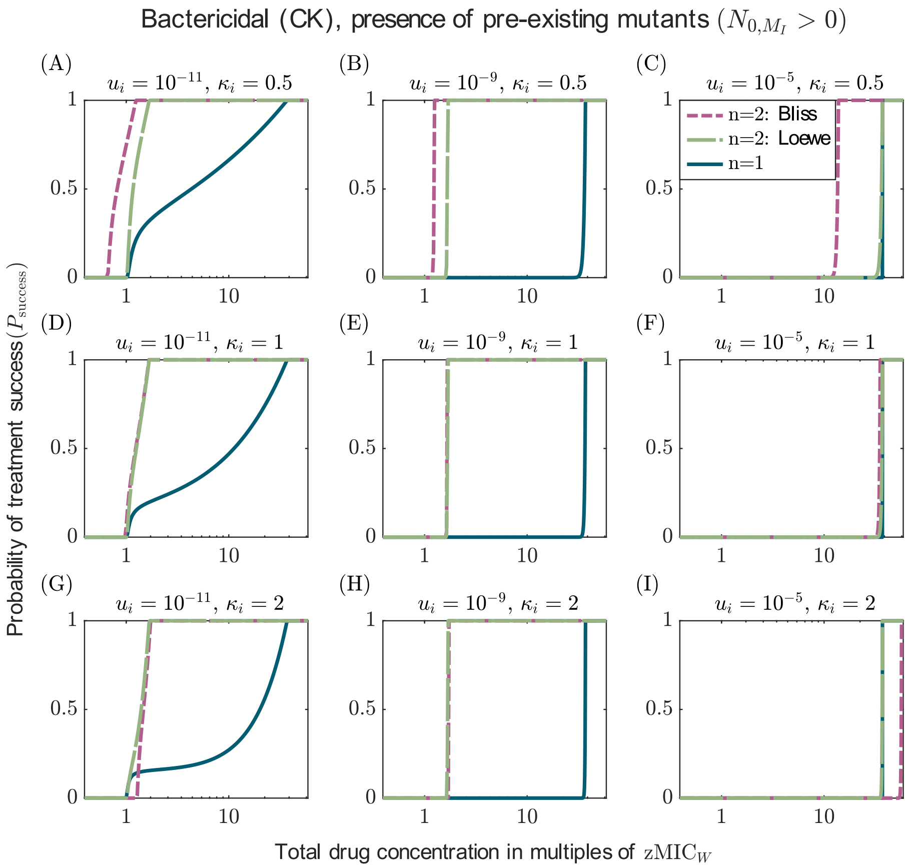


Figure S10: Comparison of the probabilities of treatment success under mono-therapy ($n = 1$) and a two-drug combination ($n = 2$) of drugs killing independent of replication (CK), when mutants pre-exist at mutation-selection balance. The figure corresponds to figure 3 with the difference that mutants pre-exist prior to treatment.

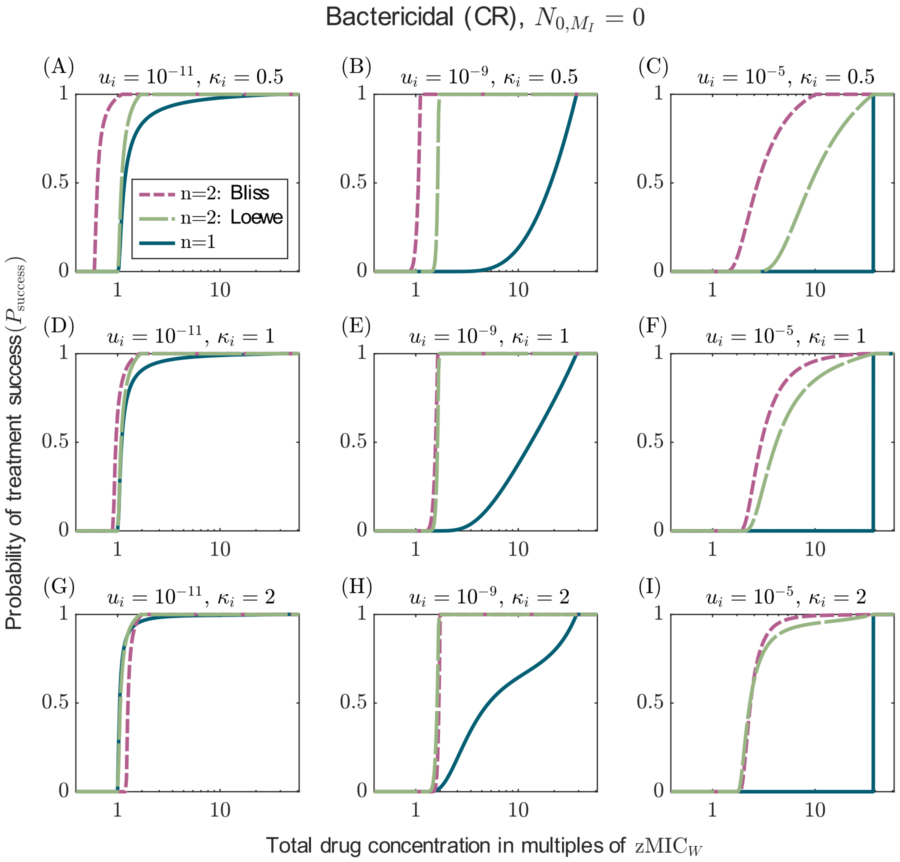


Figure S11: Comparison of the probabilities of treatment success under mono-therapy ($n = 1$) and a two-drug combination ($n = 2$) of bactericidal drugs acting during replication (CR). The figure corresponds to figure 3 with the difference that it considers CR drugs.

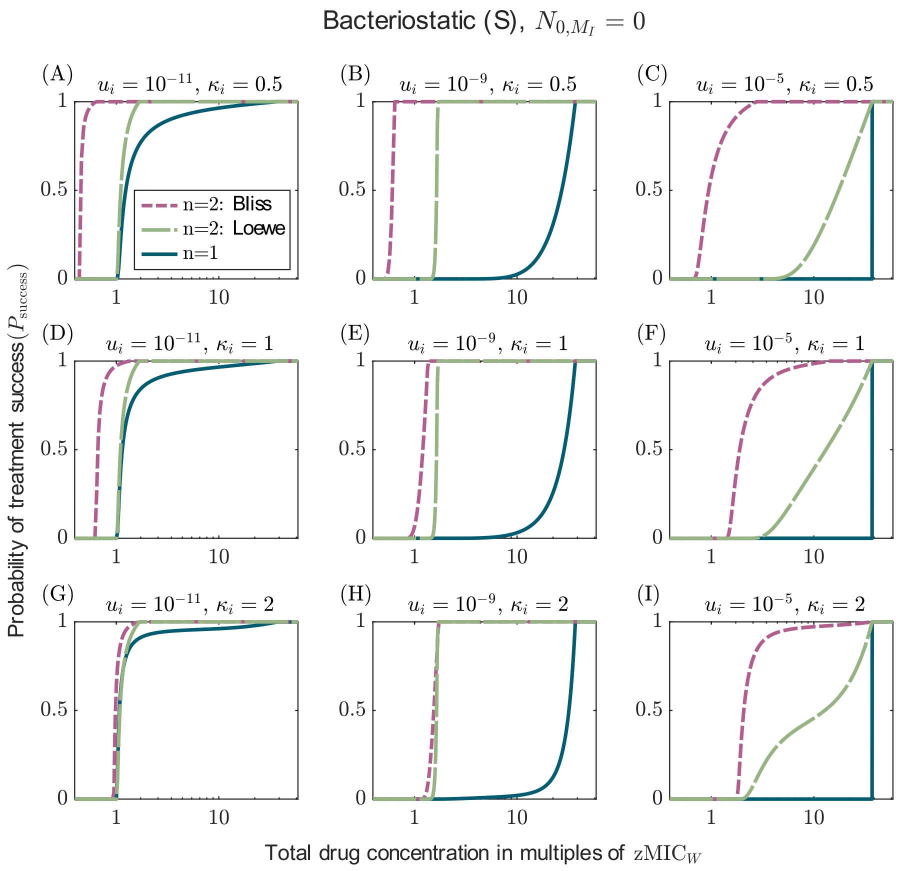


Figure S12: Comparison of the probabilities of treatment success under mono-therapy ($n = 1$) and a two-drug combination ($n = 2$) of bacteriostatic drugs (S). The figure corresponds to figure 3 with the difference that it considers bacteriostatic drugs.

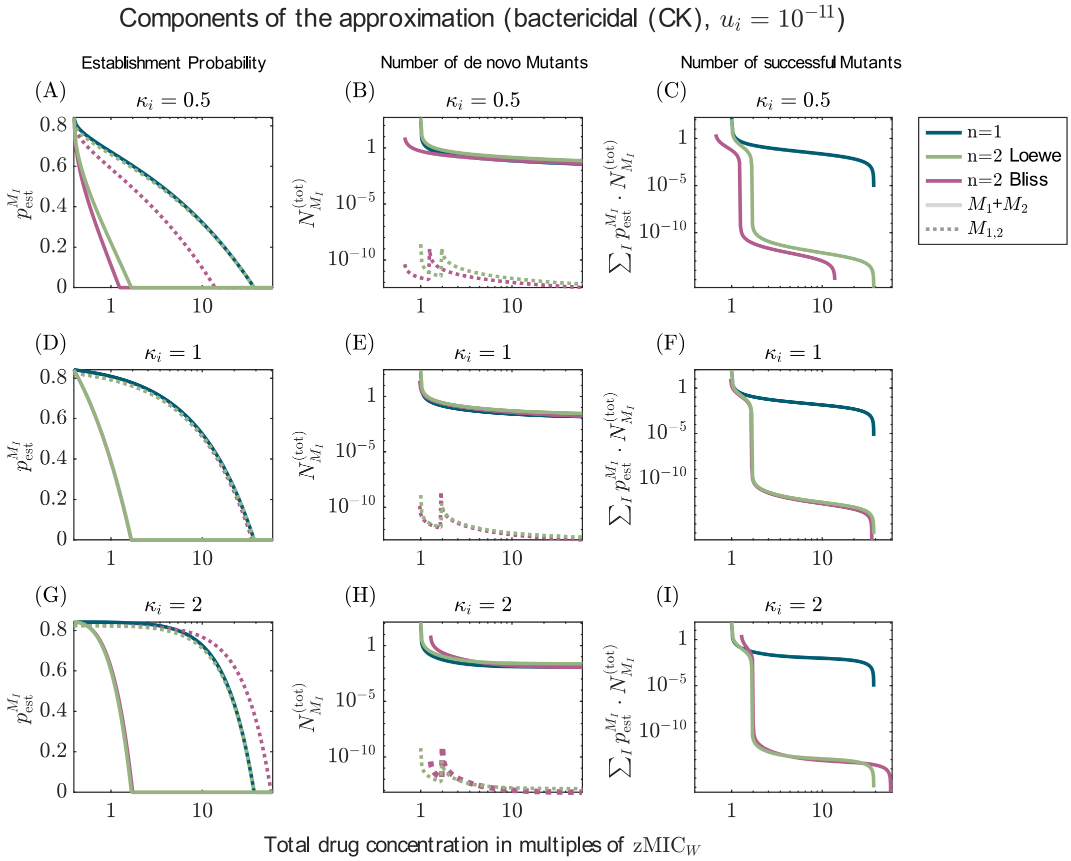


Figure S13: Components of the approximations of the probability of treatment success (equations (14), (A16), and (A21)) for mono-therapy and a two drug combination of drugs killing independent of replication. The figure is a supplement to figure 3. The rows differ by the Hill coefficient of the drugs ($\kappa = 0.5$, $\kappa = 1$, $\kappa = 2$ from top to bottom). The mutation probability is $u_i = 10^{-11}$.

#### S4 Combinations of drugs with different modes of action at a 50:50 ratio

The figures displayed in this section extend the results discussed in main text section 3.3 (see figures 4 and 5)

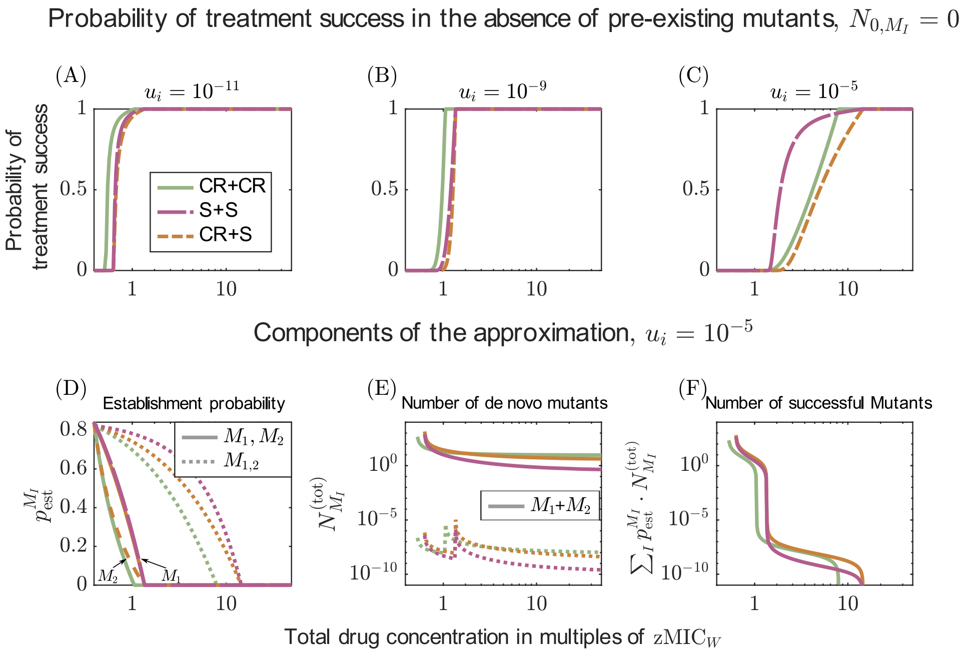


Figure S14: Comparison of the probabilities of treatment success for combinations of drugs with either the same (CR+CR and S+S) or different modes of action (S+CR). The figure corresponds to figure 4 with the difference that the drugs are of modes S and CR (instead of S and CK).

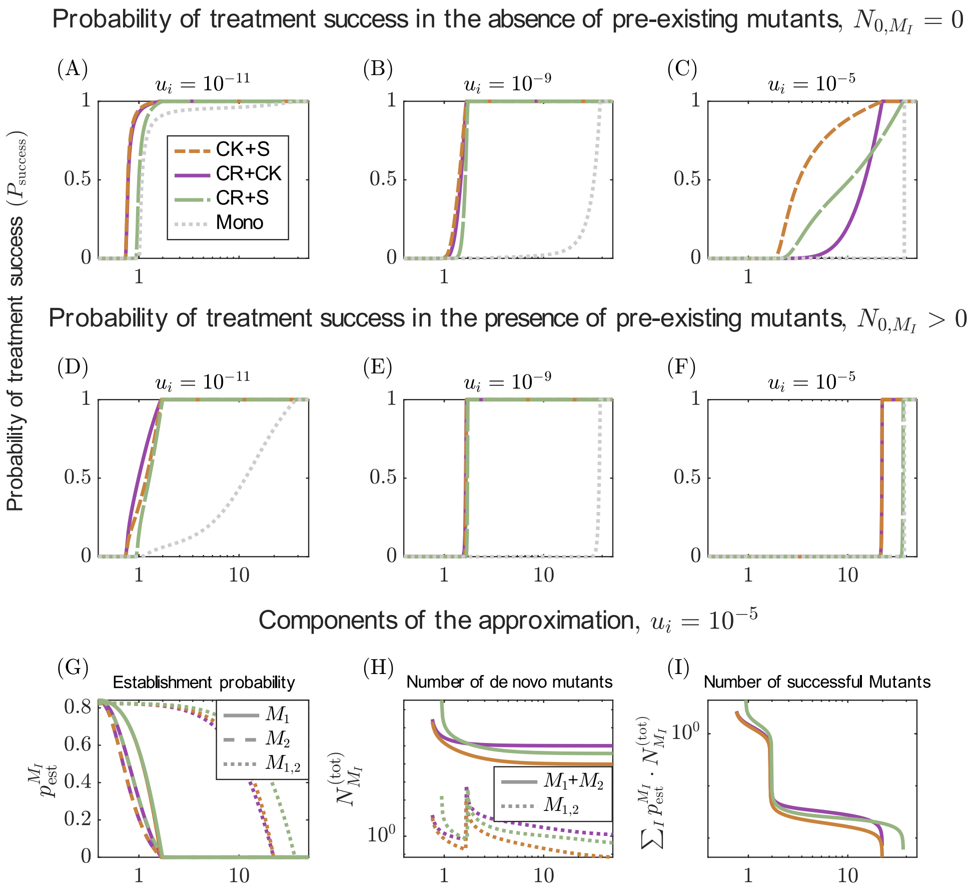


Figure S15: Comparison of the probabilities of treatment success for various combinations of drugs with different modes of action and for the best mono-therapy. The figure corresponds to figure 5 with the difference that $\kappa_i = 2$.

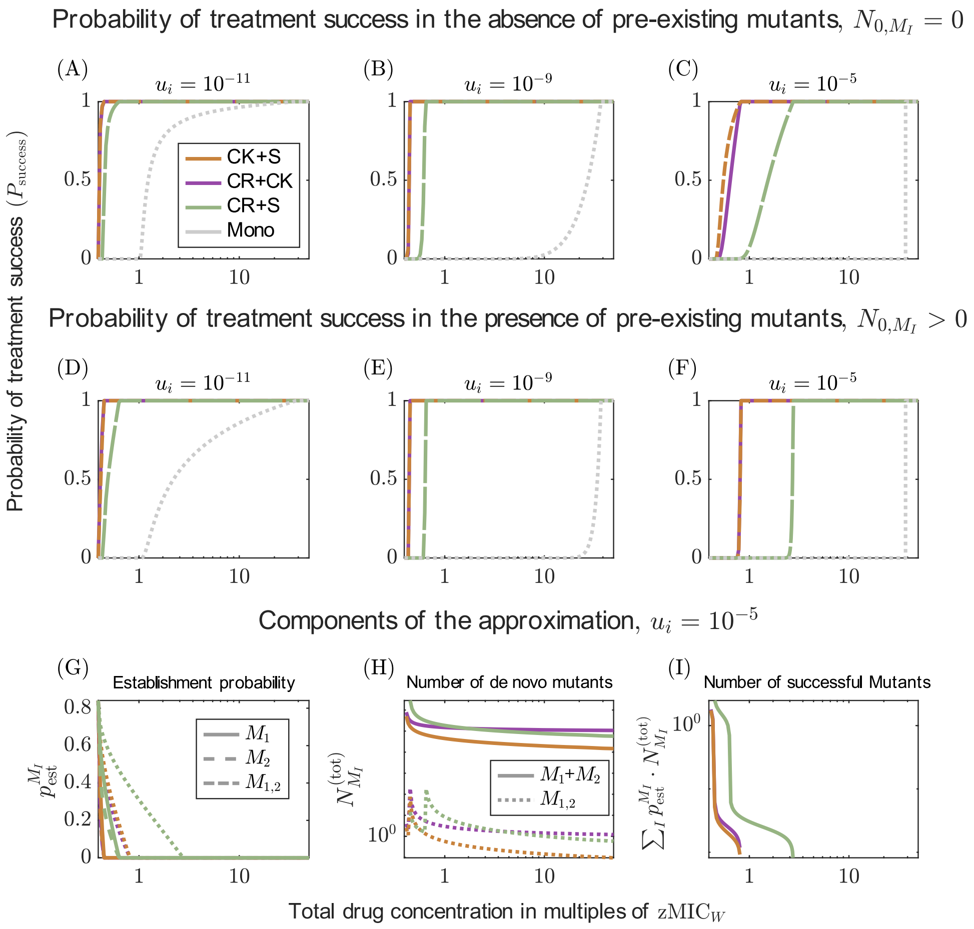


Figure S16: Comparison of the probabilities of treatment success for various combinations of drugs with different modes of action and for the best mono-therapy. The figure corresponds to figure 5 with the difference that $\kappa_i = 0.5$.

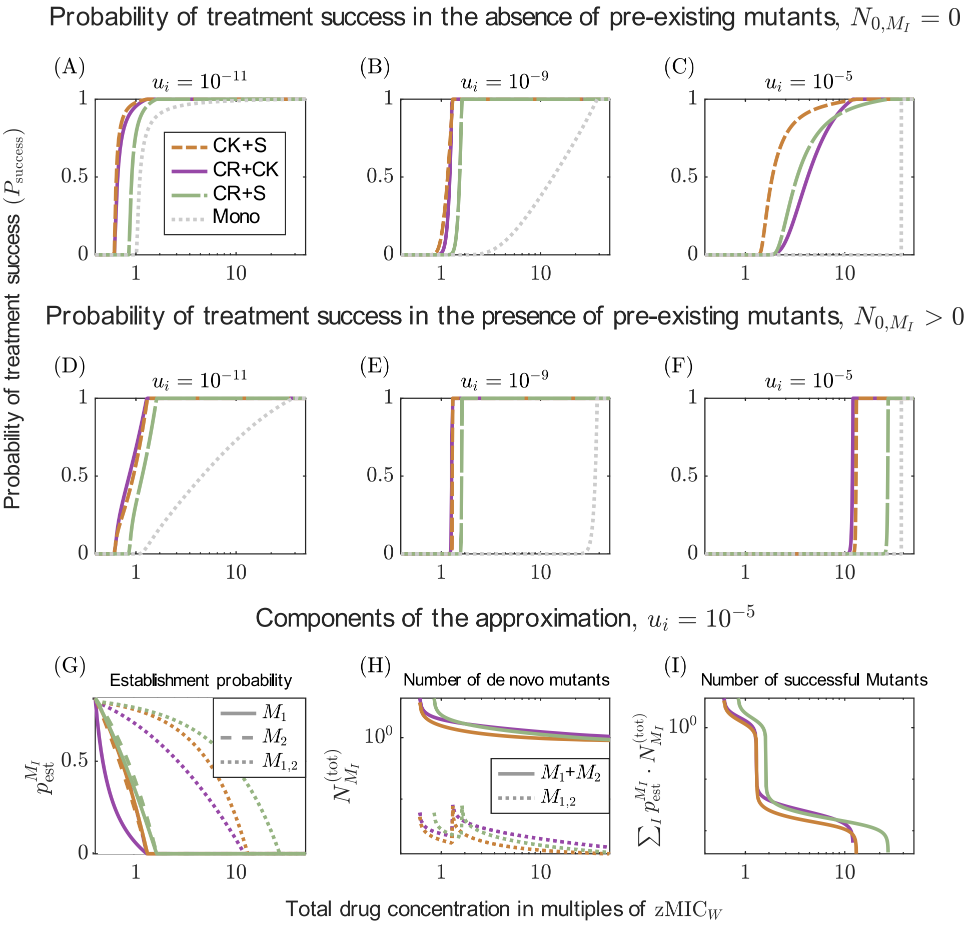


Figure S17: Comparison of the probabilities of treatment success for various combinations of drugs with different modes of action and for the best mono-therapy. The figure corresponds to figure 5 with the difference that the first drug has a larger maximum effect ($\psi_{\min}$). Here, we have for the first drug, $\psi_{\min} = -0.73^{-1}$ (CK, CR, CR); for the second drug (S,CK,S), $\psi_{\min} = -0.065^{-1}$ as before.

#### S5 Additional figures for the comparison of two-drug combinations at different drug ratios

In main text section 3.4 we displayed the comparison of treatments administered at different drug ratios for a few specific examples (see figures 6). Here, we display the comparison for more drug combinations as indicated by the the header of each figure.

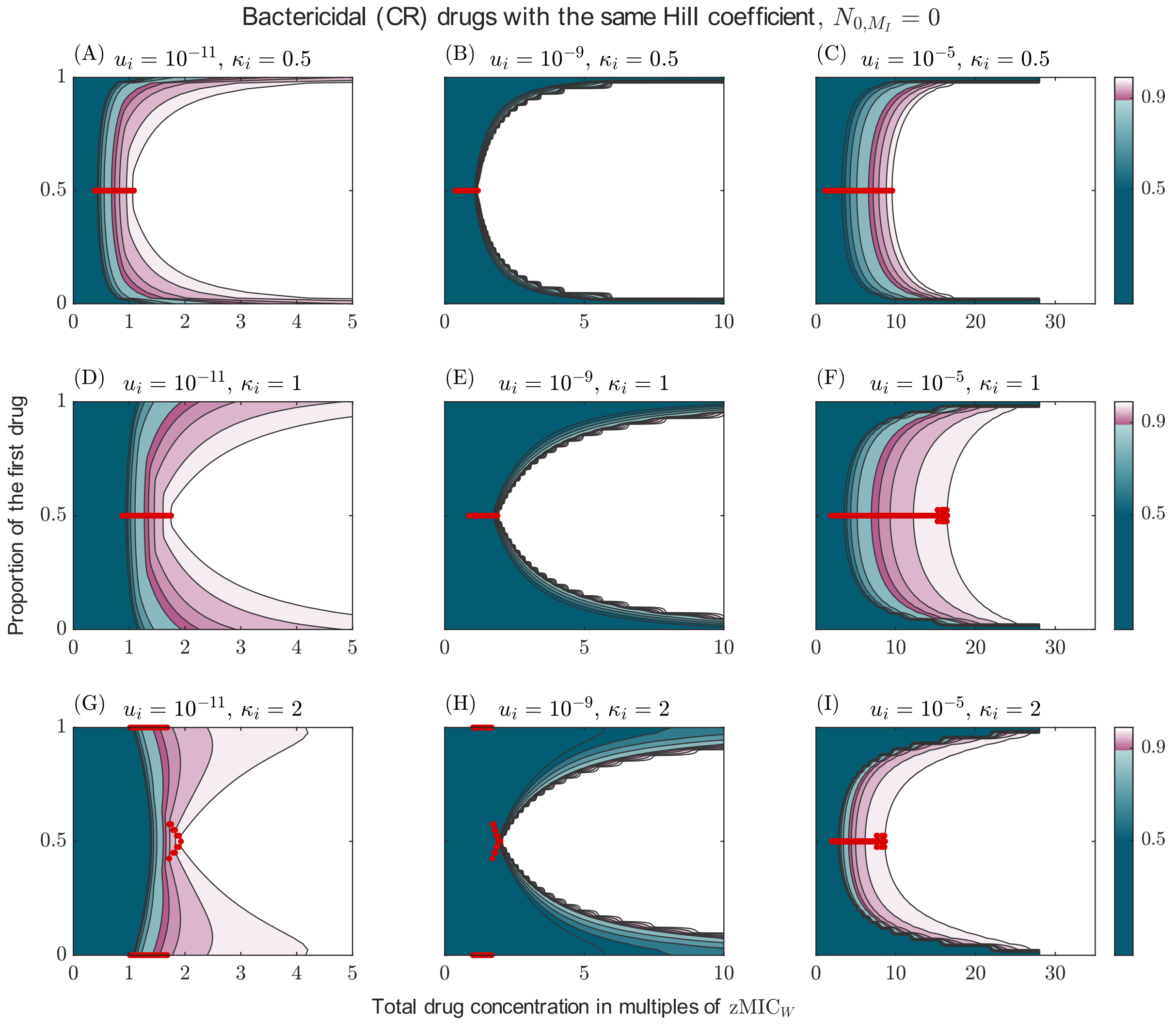


Figure S18: Probability of treatment success for combinations of two drugs administered at unequal ratios. This figure corresponds to figure 6 with the difference that it only shows combinations of bactericidal (CR) drugs ($\psi_{\min} = -0.73h^{-1}$) with the same dose-response curve, varying in steepness (rows).

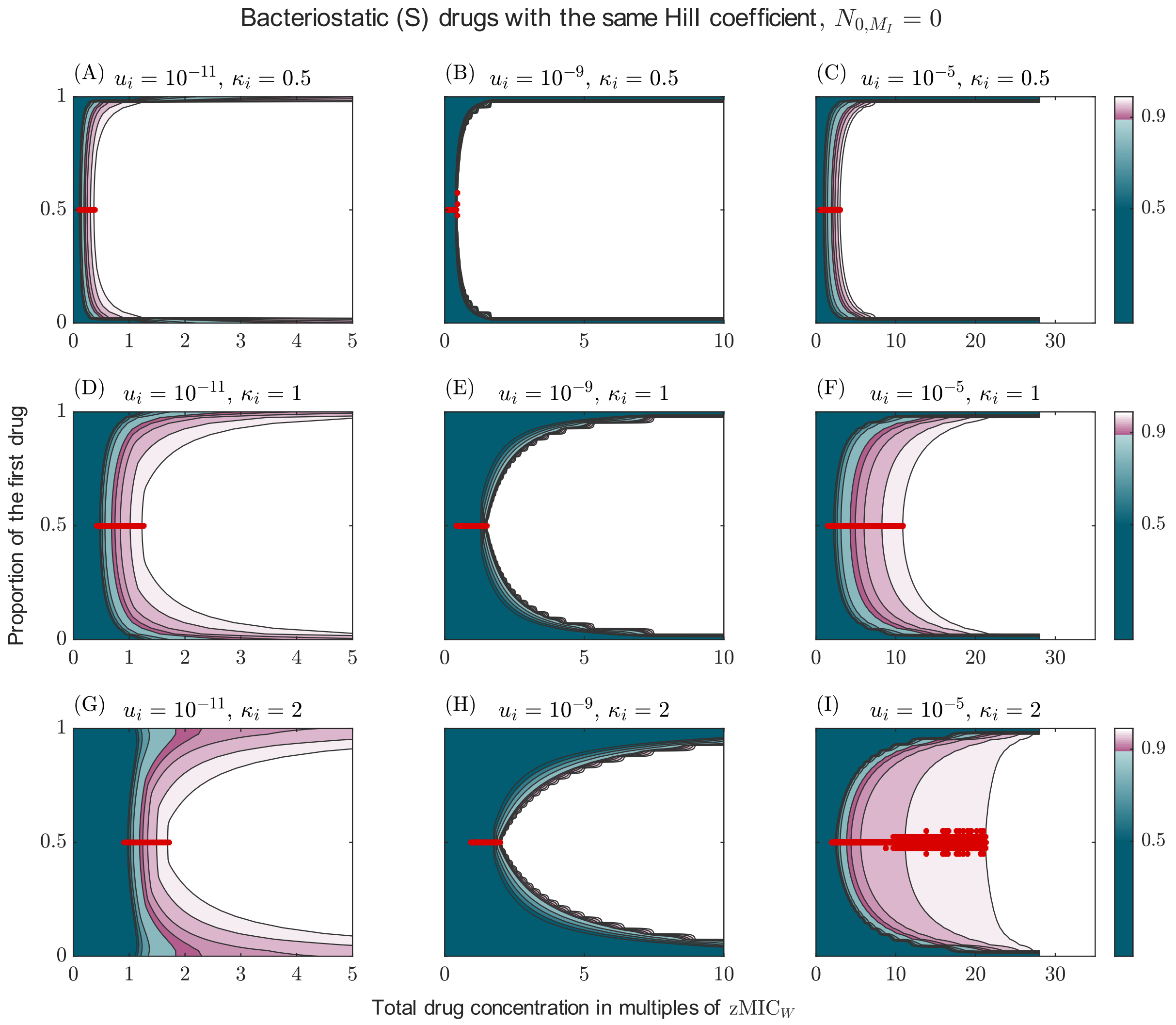


Figure S19: Probability of treatment success for combinations of two drugs administered at unequal ratios. This figure corresponds to figure 6 with the difference that it only shows combinations of bacteriostatic (S) drugs ($\psi_{\min} = -0.065h^{-1}$) with the same dose-response curve, varying in steepness (rows).

Bactericidal (CK) drugs with the same Hill coefficient, presence of pre-existing mutants $(N_{0,M_I} > 0)$

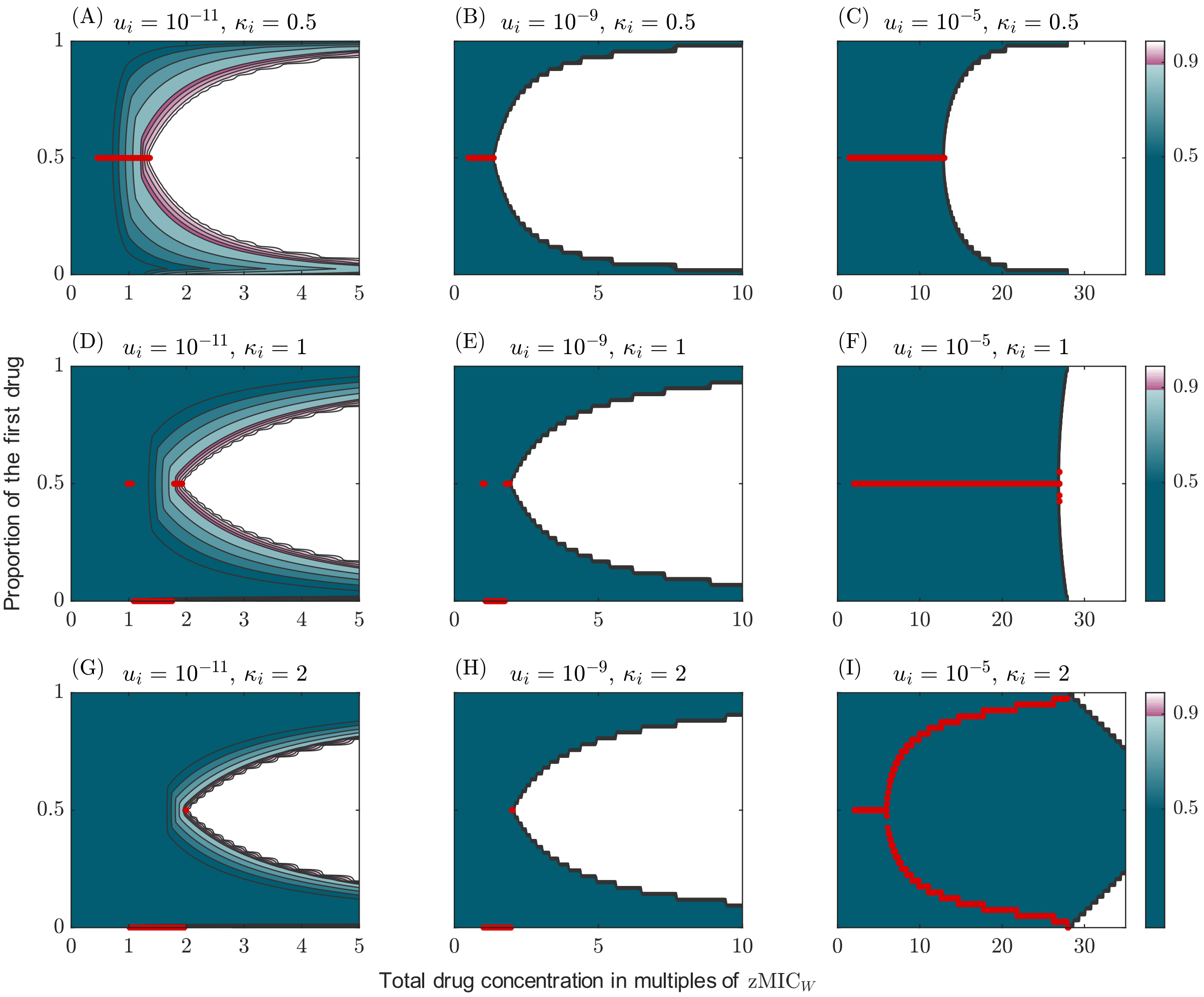


Figure S20: Probability of treatment success for combinations of two drugs administered at unequal ratios. This figure corresponds to figure 6 with the difference that it only shows combinations of bactericidal (CK) drugs ($\psi_{\min} = -6h^{-1}$) with the same dose-response curve, varying in steepness (rows) under the consideration of pre-existing mutants.

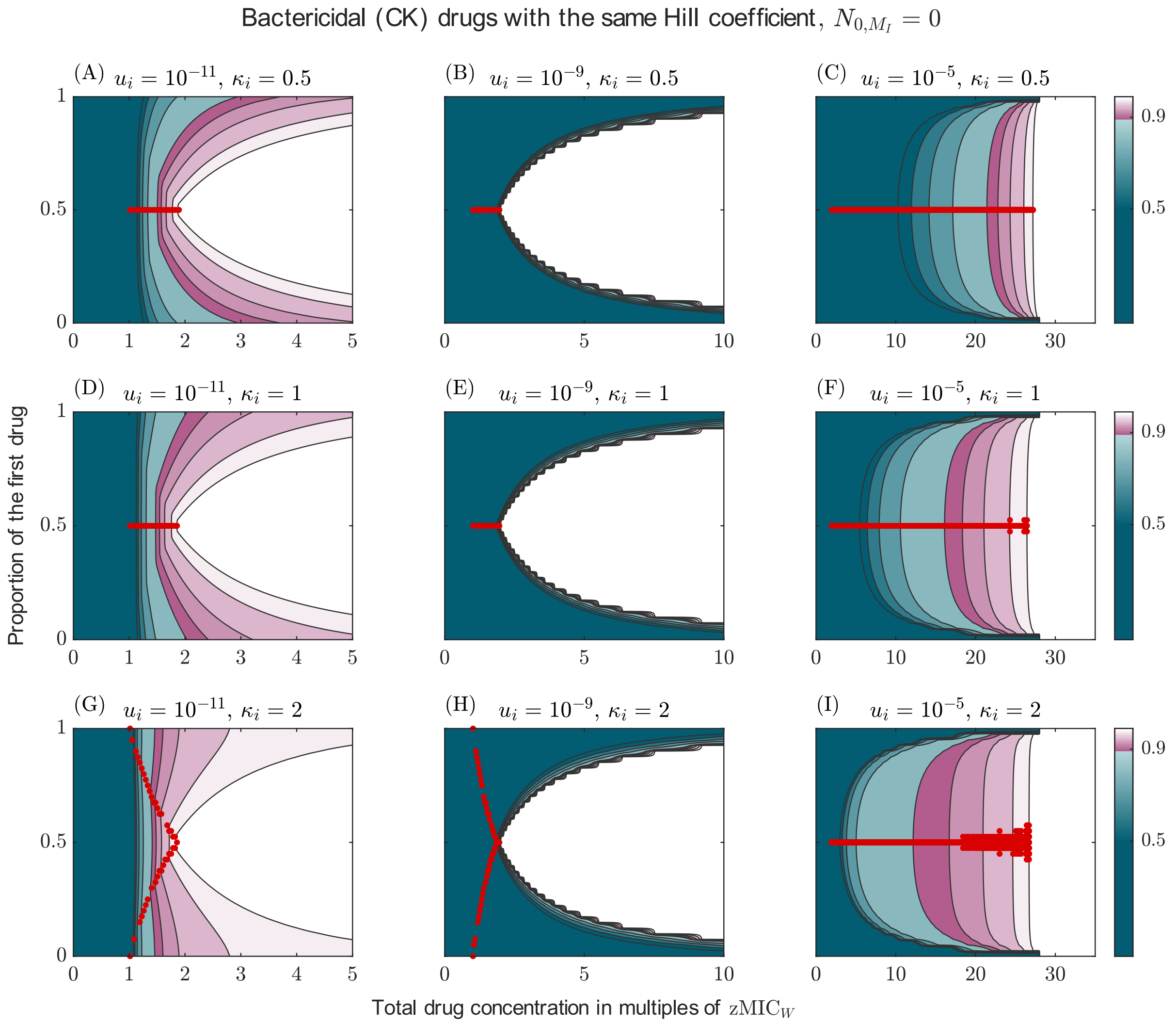


Figure S21: Probability of treatment success for combinations of two drugs administered at unequal ratios. This figure corresponds to figure 6 with the difference that it only shows combinations of bactericidal (CK) drugs ($\psi_{\min} = -6h^{-1}$) with the same dose-response curve, varying in steepness (rows), combined under Loewe additivity.

### S6 Additional figures for the comparison of treatments with more than two drugs

In this section we provide additional figures for the results on combinations with more than two drugs, discussed in main text section 3.5 (see figure 7).
The results are displayed, among other cases, for combinations with up to five drugs for either bactericidal drugs (CR) or bacteriostatic drugs (S) (see figures S22 andS23). Please note that these figures show numerical errors for the treatments with large number of drugs (the probability of treatment success should be one, but has for specific concentrations lower values). To generate the data, the fixed point of the probability generating function (which is a vector as large as the number of types and hence increases with every added drug), needs to be calculated iteratively. Small numerical error can therefore become larger with every iteration. The errors are visible, even if they only occur at the eighth or ninth digit, due to the potentiation with $N_0$.

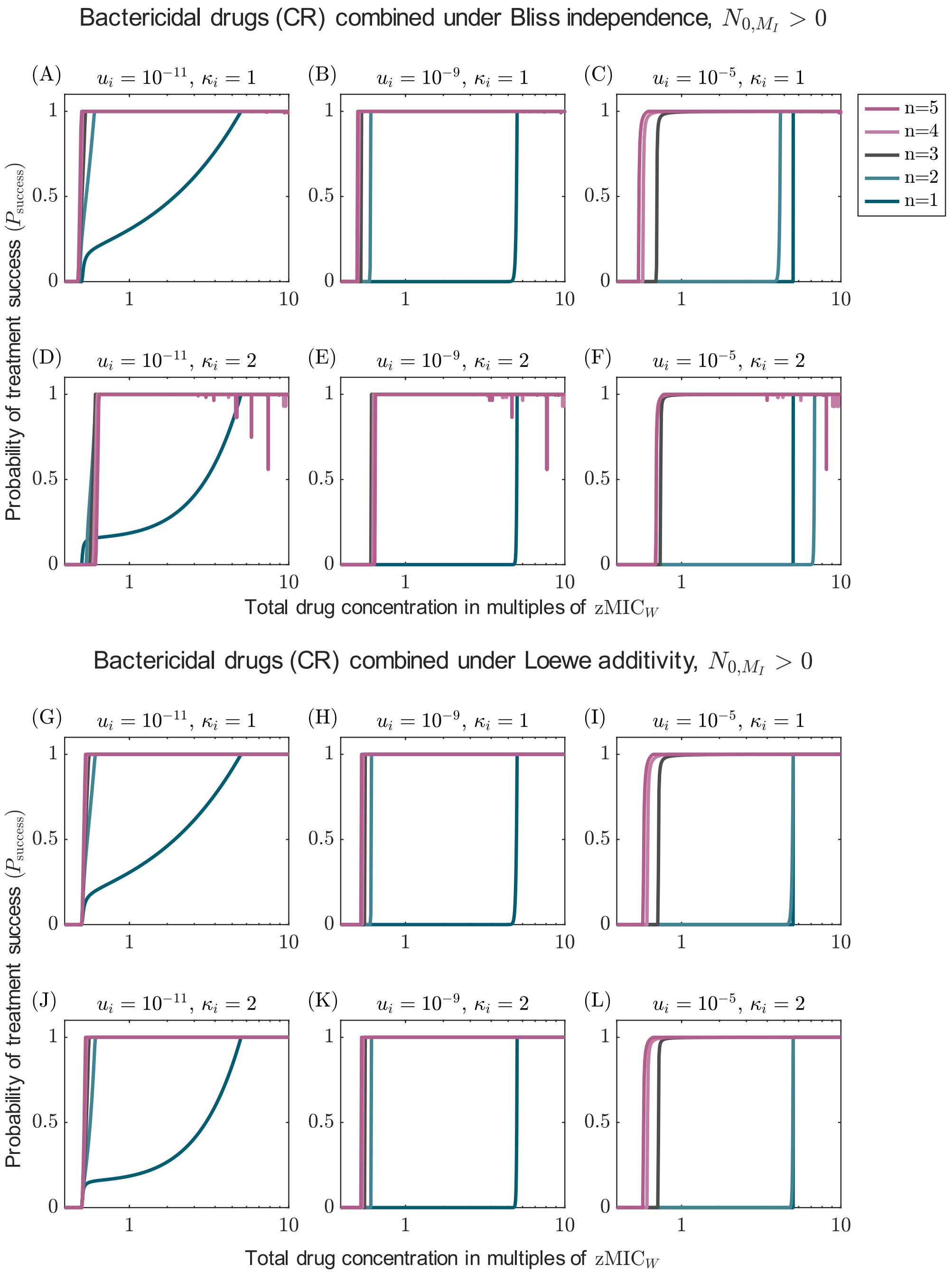


Figure S22: Probability of treatment success for combination with up to five drugs for Bliss independence and Loewe additivity for bactericidal drug acting during replication (CR), accounting for pre-existing resistance mutations. The results are displayed for different Hill coefficients (rows) as shown on the right and different mutation probabilities (columns). For large numbers of drugs, numerical artefacts appear. This figure corresponds to figure 7, except that the drugs have a different mode of action.

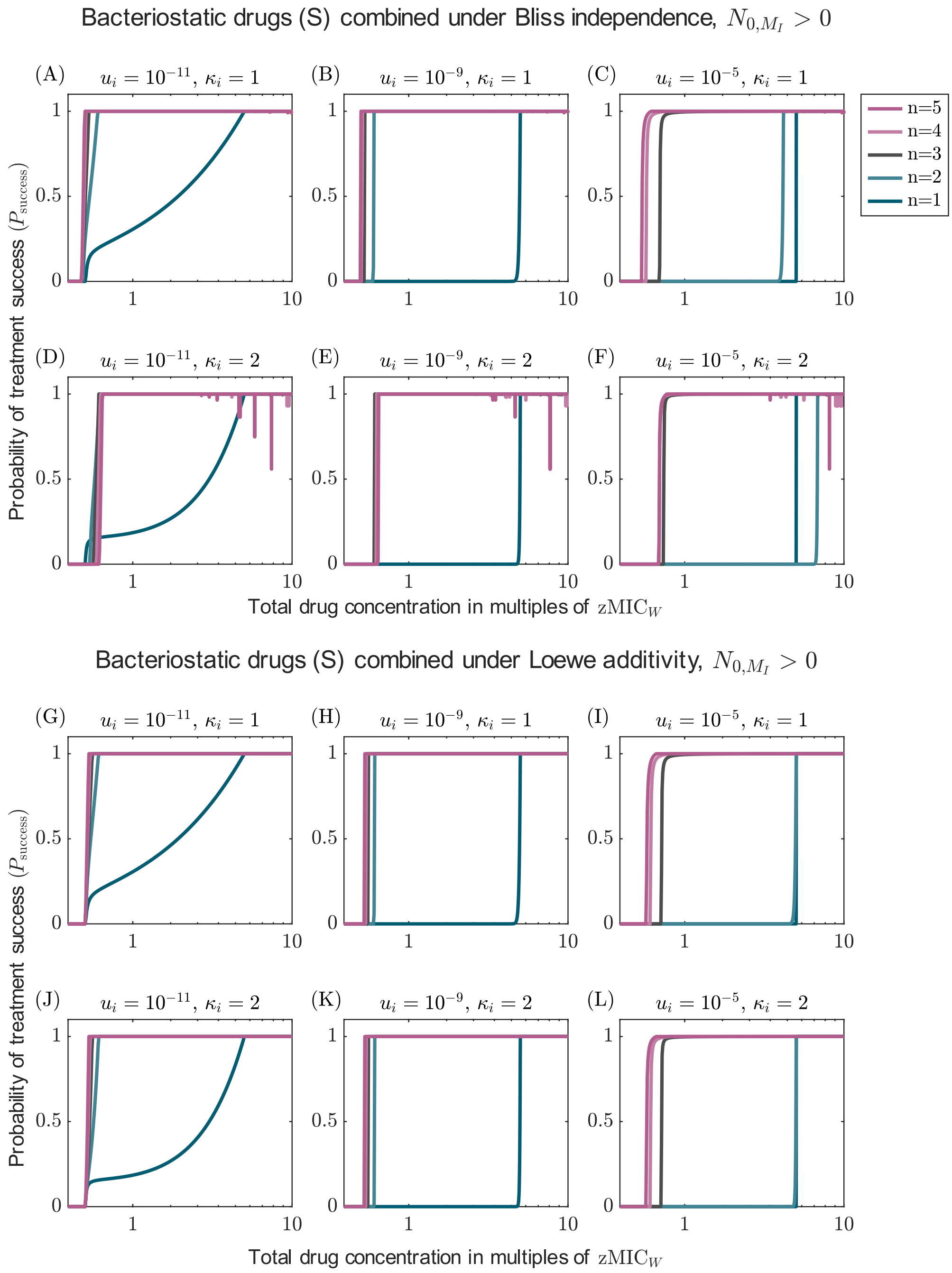


Figure S23: Probability of treatment success for combination with up to five drugs for Bliss independence and Loewe additivity for bacteriostatic drugs (S), accounting for pre-existing resistance mutations. The results are displayed for different Hill coefficients (rows) as shown on the right and different mutation probabilities (columns). For large numbers of drugs, numerical artefacts appear. This figure corresponds to figure 7, except that the drugs have a different mode of action.

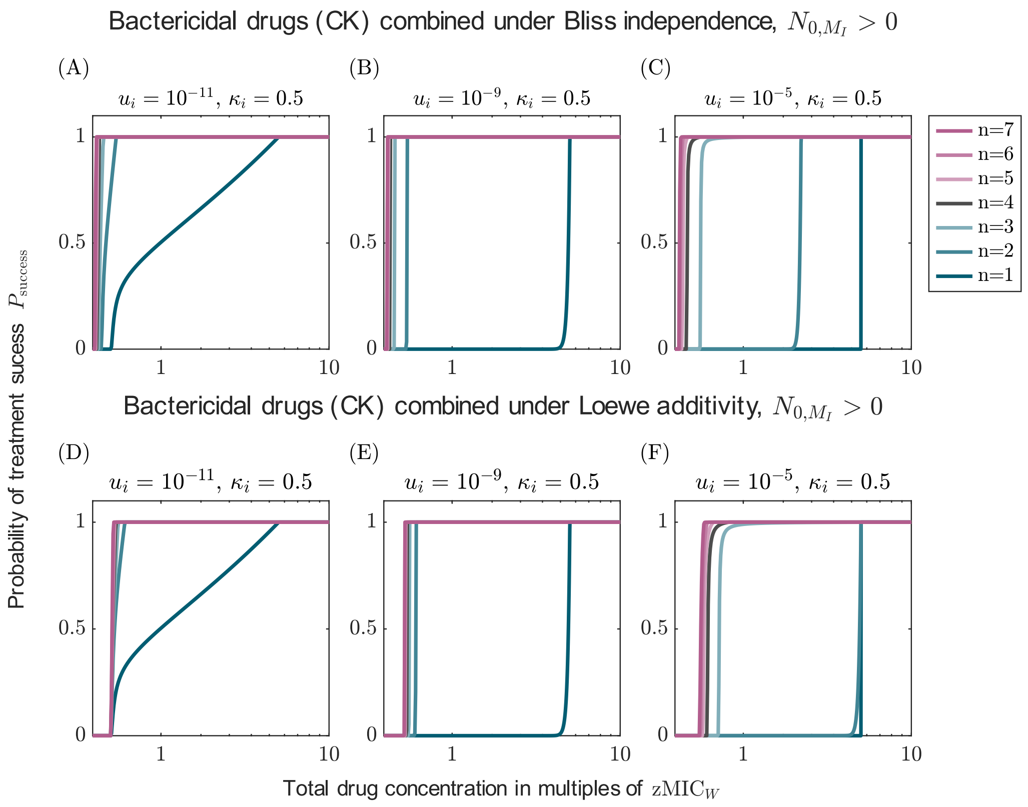


Figure S24: Probability of treatment success for combination with up to seven drugs for Bliss independence and Loewe additivity, accounting for pre-existing resistance mutations. The results are displayed for Hill coefficients $\kappa_i = 0.5$ as shown on the right and different mutation probabilities (columns). This figure corresponds to figure 7, except that the drugs have a different Hill coefficient.

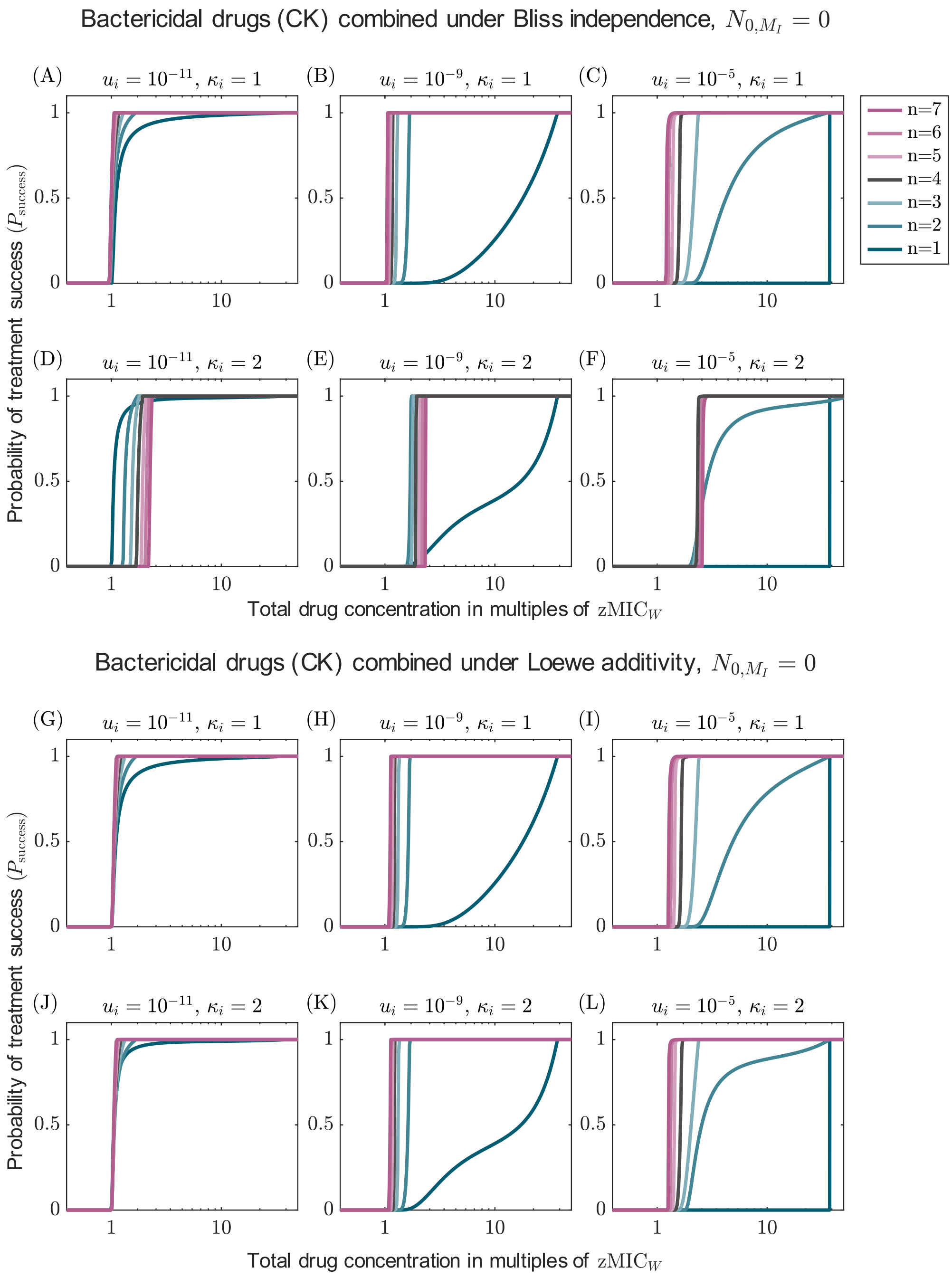


Figure S25: Probability of treatment success for combination with up to five drugs for Bliss independence and Loewe additivity in the absence of pre-existing resistance. The results are displayed for different Hill coefficients (rows) as shown on the right and different mutation probabilities (columns). This figure corresponds to figure 7, except that the resistance evolves only from *de novo* mutations.
